# Transregional astrocyte-dependent metaplasticity in the hippocampus

**DOI:** 10.64898/2026.02.10.705156

**Authors:** Shruthi Sateesh, Barbara J. Logan, Miki Suzuki, David Stellwagen, Wickliffe C. Abraham

**Author notes:** Wickliffe C. Abraham. **Email:**. **Author Contributions:** S.S. and W.C.A., designed the research. S.S., W.C.A. and D.S. obtained funding to support the research. S.S. performed the *in vitro* experiments and analyzed the data. B.L. and M.S. performed the *in vivo* experiments and analyzed the data. W.C.A. conceived the study. D.S. provided the glia-specific TNF KO mice. S.S. and W.C.A. contributed to data interpretation. S.S. wrote the manuscript, and S.S., W.C.A., B.L., M.S., and D.S. reviewed and amended the manuscript and approved the final submission. **Competing Interest Statement:** The authors declare no competing interests.

## Abstract

Learning-related synaptic plasticity is regulated by metaplasticity, which adjusts plasticity thresholds in an activity-dependent manner. We have previously described a heterodendritic form of metaplasticity whereby priming stimulation in stratum oriens (SO) inhibits subsequent long-term potentiation (LTP) in the neighboring stratum radiatum of the hippocampal CA1 region. Here, we report that this metaplasticity is, in fact, transregional, in that the SO priming stimulation also inhibits later LTP induction at dentate gyrus (DG) middle molecular layer (MML) synapses, both *in vitro* and *in vivo*. This effect operates across the hippocampal fissure since it occurs in the absence of CA3, thus highlighting a previously unappreciated reverse-direction and long-distance hippocampal crosstalk. Our findings demonstrate an essential role of astrocytes as SO priming elicited a sustained increase in the frequency of calcium (Ca^2+^) events in astrocytes in the DG MML, while the metaplasticity effect was blocked by calcium-buffering in MML astrocytes. It could be triggered by either activation of M1 muscarinic acetylcholine receptors or group II metabotropic glutamate receptors, and was critically dependent on inositol 1,4,5-trisphosphate receptor type 2 signaling. The transregional inhibition of LTP was mediated by astrocytic release of tumor necrosis factor (TNF), which likely acts in an autocrine fashion on astrocytic TNF type 1 receptors (TNFR1s). Downstream of TNF-TNFR1 signaling, the inhibition of MML LTP was mediated by the activation of GluN2B-containing N-methyl-D-aspartate receptors. Thus, a complex, bidirectional neuron-glia signaling cascade orchestrates long-distance metaplasticity across hippocampal subregions, providing a novel framework for understanding how hippocampal neuronal networks dynamically regulate plasticity thresholds across space and time.

**Significance Statement:** This study documents the existence of a transregional metaplasticity that traverses from area CA1 to the dentate gyrus across the hippocampal fissure. This reveals a novel long-distance crosstalk within the hippocampus, in addition to the classical pathways of information transfer. This form of metaplasticity entails an essential contribution by astrocytes that integrate neuronal activity and use a complex Ca^2+^-dependent neuron-glia-neuron signaling cascade to dynamically regulate LTP thresholds in the hippocampal dentate gyrus.

## Introduction

Synaptic plasticity refers to the activity-dependent and long-lasting changes in the efficacy of synaptic connections, such as long-term potentiation (LTP) and long-term depression (LTD). Such plasticity is fundamental to the ability of neural circuits to adapt and store information [1]. Moreover, it is subject to higher-order regulation such as neuromodulation and metaplasticity. Metaplasticity, defined as an activity-dependent change in the ability to induce subsequent synaptic plasticity [2], plays a critical role in optimizing neuronal network function for efficient information encoding and memory storage [3].

While canonical models of synaptic plasticity and metaplasticity often focus on localized changes within individual synapses, there is evidence that plasticity thresholds can be dynamically regulated across spatially distinct dendritic compartments. Previously, we described a unique Bienenstock, Cooper, and Munro-like (BCM) [4] form of metaplasticity in hippocampal CA1 pyramidal cells, whereby electrical “priming” stimulation delivered to afferents in the basal dendrite region of stratum oriens (SO) inhibited subsequent LTP and enhanced LTD at apical synapses located in a spatially separate dendritic compartment, stratum radiatum (SR) [5]. However, unlike the BCM model, this form of heterodendritic metaplasticity operates independently of postsynaptic action potential firing [5]. Further, we have demonstrated that this form of metaplasticity is highly pathway-specific within the CA1 region of the hippocampus, affecting synapses in SR, but not in SO or stratum lacunosum-moleculare (SLM), in a tumor necrosis factor (TNF)-dependent fashion [6]. This synapse-specific long-range interaction between widely separated dendritic regions raises interesting and important questions as to the mechanisms by which this effect is mediated across space and time.

We have previously shown that the long-range inhibition of SR LTP is blocked by an inhibitor of the enzyme converting ATP to adenosine and mimicked by administration of an adenosine 2B receptor (A2BR) agonist, both in vitro and in vivo [7]. Interestingly, the effect was also blocked by the gap junction inhibitors carbenoxolone and meclofenamic acid, and by a peptide antagonist of connexin-43, a key component of astrocytic gap junctions and hemichannels [8]. The requirement of inositol triphosphate receptor-dependent (IP_3_R_2_) Ca^2+^ release in astrocytes underscored the critical role of astrocytes in mediating the heterodendritic metaplasticity effect on SR LTP [5, 7]. Given that the astrocytic network forms a functional syncytium capable of crossing physical barriers, such as the hippocampal fissure [9], we hypothesized that this metaplasticity could potentially extend beyond CA1 to influence synaptic plasticity elsewhere in the hippocampus. Specifically, we investigated whether SO synaptic priming regulates LTP through astrocyte activation even in the dentate gyrus (DG), a region adjacent to CA1 but separated from it by the hippocampal fissure, and which hosts a very different type of principal cell, the granule cell. We found that the metaplasticity effect indeed spreads transregionally. Thus, priming stimulation in the SO of CA1 strikingly inhibited subsequent LTP in the middle molecular layer (MML) of the DG. Our findings revealed a complex, multi-step signaling cascade that orchestrates the transregional effect. We provide direct evidence that astrocytes in the MML of the DG respond to SO priming stimulation with increased astrocytic Ca^2+^ events both immediately following and 30 minutes post-priming. We confirmed that Ca^2+^ signaling via IP_3_ type 2 receptor-mediated Ca^2+^ release from the endoplasmic reticulum is essential for the SO priming-induced inhibition of LTP in MML. This transregional metaplasticity could be triggered by either activation of muscarinic acetylcholine receptors (mAChRs) or group II metabotropic glutamate receptors (mGluRs), and then by release of TNF from astrocytes, but not microglia. The TNF appeared to subsequently trigger astrocytic glutamate release, activating ifenprodil-sensitive extrasynaptic GluN2B-containing N-methyl-D-aspartate receptors (NMDARs) in the nearby synapses in the DG MML [10-12]. These findings reveal a complex, bidirectional neuron-glia signaling cascade that orchestrates long-distance metaplasticity across hippocampal subregions, challenging traditional views of localized synaptic plasticity regulation.

## Results

### Electrical priming in SO inhibits subsequent LTP in DG MML *in vitro* and *in vivo*

Previously, we demonstrated synapse-specific heterodendritic metaplasticity in area CA1 from stimulating in SO [6]. Here, we investigated whether this metaplasticity effect extended into the MML of the dentate gyrus of male rats. Similar to our previous observations in SR of CA1, HFS priming (600 pulses) in SO produced homosynaptic LTP in the SO pathway of CA1 slices (**Fig. S3A**) without affecting baseline responses in the MML. Importantly, the SO HFS priming strongly inhibited subsequent LTP that was induced 30 minutes later at MPP synapses in the MML (**Fig. 1A**).

**Figure 1.**
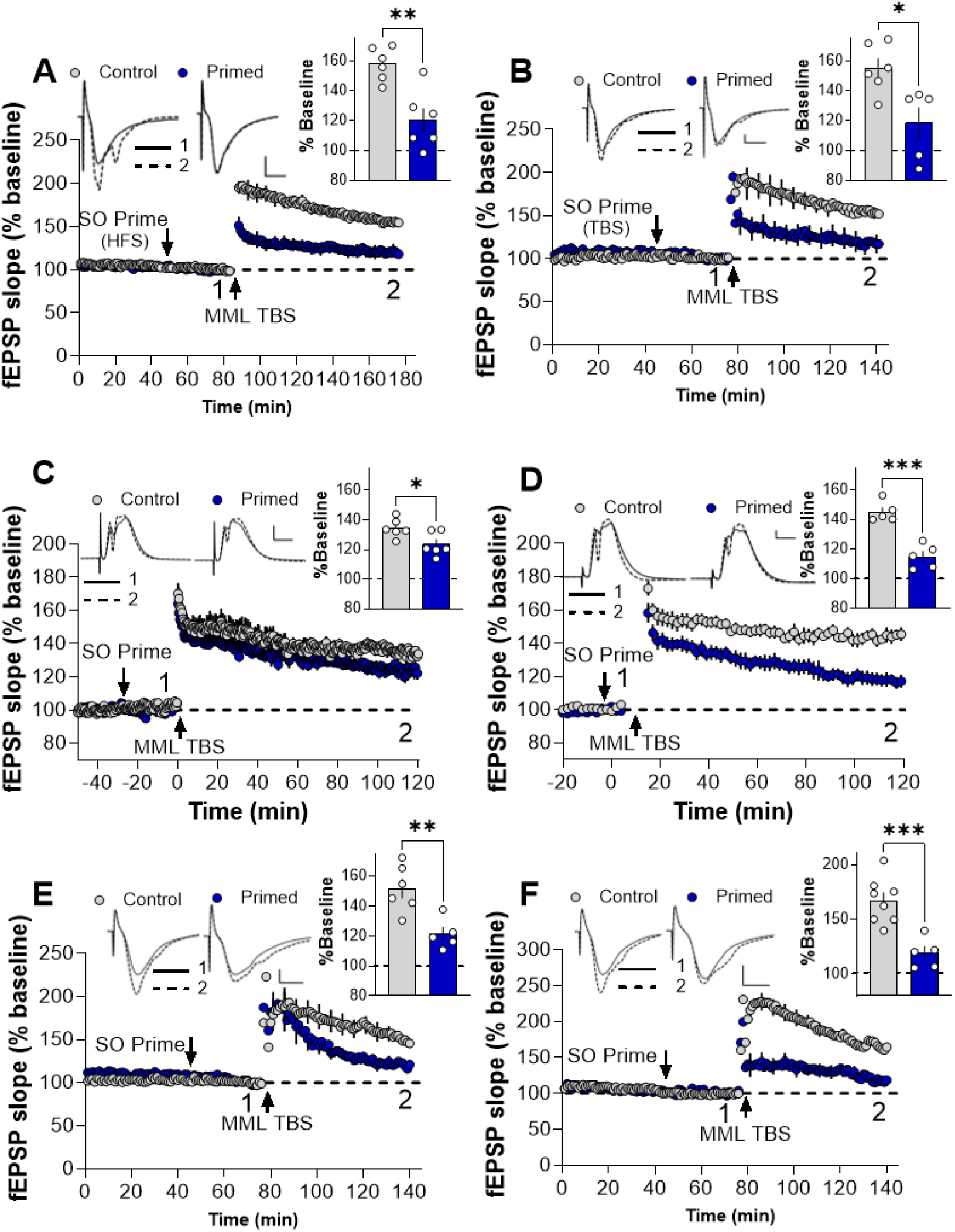
Electrical priming in SO inhibits subsequent LTP in DG MML *in vitro* and *in vivo* and is independent of sex. **(A)** HFS priming in SO *in vitro* inhibited MML LTP induced 30 min later (Control non-primed: 157.8 ± 4.5%, *n* = 6; Primed: 120.2 ± 8%, *n* = 6, *t*_*(10)*_ = 4.1, *p* = 0.0021). **(B)** TBS priming in SO *in vitro* inhibited MML LTP induced 30 min later (Control non-primed: 155 ± 6.7%, *n* = 6; Primed: 118.2 ± 10.7 %, *n* = 5, *t*_*(9)*_ = 3.0, *p* = 0.0142). TBS priming in SO *in vivo* inhibited MML LTP that was induced **(C)** 30 min post-priming (Control non-primed: 134.5 ± 2.6 %, *n* = 6; Primed: 123.5 ± 3.3 %, *n* = 6, *t*_*(10)*_ = 2.6, *p* = 0.0250), and **(D)** 5 min post-priming (Control: 145.1 ± 3%, *n* = 5; Primed: 115 ± 3.6 %, *n* = 5, *t*_*(8)*_ = 6.4, *p* = 0.0002) in anesthetized rats. **(E**,**F)** TBS priming in SO inhibited MML LTP induced 30 min later in the mouse hippocampus of both **(E)** males (Control non-primed: 151.7 ± 6.4 %, *n* = 6; Primed: 121.3 ± 4.5 %, *n* = 5, *t*_*(9)*_ =3.7, *p* = 0.0046) and **(F)** females (Control non-primed: 167 ± 7.3 %, *n* = 7; Primed: 118.8 ± 6.2 %, *n* = 5, *t*_*(11)*_ = 4.5, *p* = 0.0008). Arrows indicate the timing of SO priming and MML LTP induction. ***Insets:*** Representative waveforms are an average of 10 synaptic responses prior to LTP induction (**1**) and at the conclusion of the experiment (**2**). Bar graph summarizes LTP across individual slices. Scale bars: 0.5 mV, 5 ms (**A-B**), 3 mV, 5 ms (**C**), 5 mV, 5 ms (**D**), 1 mV, 5 ms (**E-F**). All data presented as mean ± SEM; ∗, *p* < 0.05; ∗∗, *p* < 0.01; ∗∗∗, *p* < 0.001.

We have previously shown that TBS priming with fewer pulses can also generate the equivalent priming effect in CA1 SR [6]. Likewise, we found here that priming with the TBS pattern of activity (100 pulses) significantly inhibited MML LTP compared to control slices (**Fig. 1B**), while also generating homosynaptic LTP in the SO priming pathway (**Fig. S3B**). Taken together, these data demonstrate the correspondence of the priming-mediated LTP inhibition in DG with that previously observed in CA1.

To confirm the physiological relevance of these findings, we extended our investigation to an *in vivo* model. Consistent with the inhibitory effects on MML LTP observed *in vitro*, TBS priming in SO significantly impaired MML LTP induced 30 min later in anesthetized rats (**Fig. 1C**). To further assess the robustness of the SO TBS priming-mediated inhibition of MML LTP, we reduced the time interval between the priming and conditioning stimuli to 5 min. Even with this reduced interval, the priming stimulation, which induced substantial homosynaptic LTP in the SO (**Fig. S4A**), strongly inhibited MML LTP both *in vivo* (**Fig. 1D**) and *in vitro* (**Fig. S4B**).

We next investigated whether the transregional metaplasticity was species- or sex-dependent. As seen above for male rats, TBS priming in SO robustly inhibited subsequent LTP in the MML of both male (**Fig. 1E**) and female mice (**Fig. 1F**), and to a similar extent. As expected, the TBS pattern of activity in the SO priming pathway resulted in substantial potentiation as measured 30 min post-priming in both males (**Fig. S5A**) and females (**Fig. S5B**). Interestingly, unlike for the rat slices with CA3 removed, the SO LTP in these experiments with intact CA3 underwent complete depotentiation following the conditioning stimulation in MML, such that it was no longer significantly different from non-primed controls in both sexes (**Fig. S5A**,**B**).

### Intercellular signaling mediating transregional metaplasticity

Having established that priming can regulate synaptic plasticity across significant distances between sub-regions containing different cell types—even crossing the hippocampal fissure—we next investigated the contributions of the astrocytic network. To directly confirm its activation during the period of metaplasticity, we undertook Ca^2+^ imaging in DG astrocytes located in the MML in the presence and absence of electrical priming in SO. For this, we used cre-inducible astrocyte-specific Ald1L1/cre-ERT2^+/-^ mice crossed with Ai148 GCaMP6f (TIT2L-GC6f-ICL-tTA2)^+/-^, which were administered tamoxifen and used one to two weeks post-injections (**Fig. 2A**). We observed a significant increase in the number of astrocytic Ca^2+^ events immediately following priming stimulation (0-10 minutes) and an ongoing enhancement in the frequency of these Ca^2+^ events (20-30 minutes) post-priming (**Fig. 2B, SV1**) compared to non-primed controls which showed a gradual decline in Ca^2+^ event frequency over time (**Fig. 2B**).

**Figure 2.**
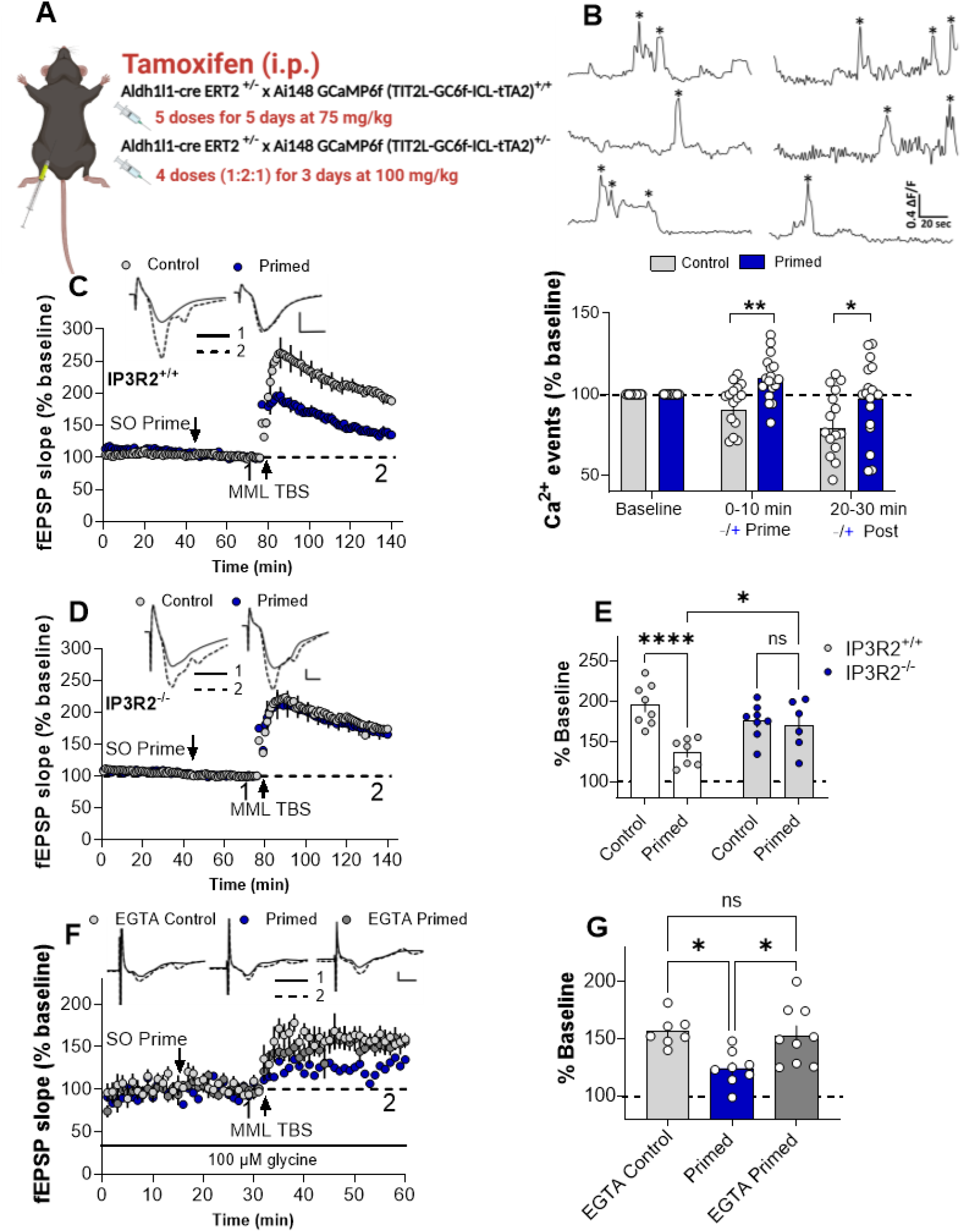
Transregional metaplasticity in the hippocampus requires astrocytic Ca^2+^. **(A)** Schematic showing the tamoxifen administration for the Aldh1l1-Cre/ERT2^+/-^ x homozygote and heterozygote Ai148 GCaMP6f transgenic mice. **(B)** Representative traces of Ca^2+^ fluorescence intensity (ΔF/F) in ML astrocytes over time. Asterisks indicate calcium transients detected in a defined ROI by AQuA software and sample video file. Two-way ANOVA indicated an overall main effect of priming (*F*_*(1, 30)*_ = 15.35, *p* = 0.0005) as well as a significant change in the frequency Ca^2+^ events across time (*F*_*(1, 34)*_ = 5.612, *p* = 0.0237). Post-hoc Uncorrected Fishers LSD test revealed in a significant increase in the number of astrocytic Ca^2+^ events measured 0 - 10 min post-priming (% baseline: Control: 90.1 ± 5.0%, *n* = 16; Primed: 110.0 ± 3.2%, *n =* 18, *p* = 0.0069) and an ongoing increase of these Ca^2+^ events as measured 20 - 30 min post-priming (Control: 79.3 ± 5.9%, *n* =16; Primed: 96.9 ± 5.5%, *n* = 18, *p* = 0.0130). **(C-E)** Transregional metaplasticity required astrocytic IP_3_R2 receptors. Two-way ANOVA indicated a significant priming effect (*F*_*(1, 25)*_ = 12.78, *p* = 0.0015). Importantly, while SO TBS priming readily inhibited LTP in the **(C**,**E)** IP_3_R2^+/+^ mice (Control non-primed: 196.1 ± 9.1%, *n* = 8; Primed: 137.4 ± 6.4%, *n* = 7, post hoc Uncorrected Fishers LSD test, *p* <0.0001), this priming effect was absent in the **(D**,**E)** IP_3_R2^-/-^ mice (Control: 175.8 ± 7.6%, *n* = 8; Primed: 170.5 ± 12.7%, *n* = 6, *p* = 0.69). **(F-G)** Transregional metaplasticity required astrocytic Ca^2+^ activity in the MML. One-way ANOVA revealed a significant main effect of treatment group (*F*_*(2, 21)*_ = 6.5, *p* = 0.0065). MML LTP was inhibited by SO priming in the absence of the Ca^2+^ clamp when compared to non-primed EGTA controls (EGTA Control non-primed: 156.7 ± 5.0%, *n* = 7; Primed: 124.3 ± 5.3%, *n* = 8, post hoc Tukey’s test, *p* = 0.0113). MML LTP post-priming was restored to control levels in the presence of the Ca^2+^ clamp (EGTA Primed: 152.8 ± 8.6%, *n* = 9, *p* = 0.92) compared to in the absence of the Ca^2+^ clamp (*p* = 0.0176). **(E**,**G)** Bar graphs summarize LTP across individual slices for the experiments described above. Arrows indicate the timing of SO priming and MML LTP induction. *Inset:* Representative waveforms are an average of 10 synaptic responses prior to LTP induction **(1)** and at the conclusion of the experiment (**2**). Scale bars: 1 mV, 5 ms (**C**), 0.5 mV, 2 ms (**D**), 2 mV, 2 ms (**F**). All data presented as mean ± SEM; ∗, *p* < 0.05; ∗∗, *p* < 0.01; ∗∗∗∗, *p* < 0.001.

The fact that the frequency of astrocytic Ca^2+^ events was significantly increased post-priming indicates a likely role for tripartite synapse signaling in mediating the metaplasticity effect. Accordingly, we tested the effect of SO electrical priming on MML LTP in IP_3_R2^-/-^ mice, since IP_3_R2 activation is a primary generator of astrocytic Ca^2+^ signaling, which in turn triggers the activity-dependent release of gliotransmitters [22, 23]. First, IO curves were assessed at MML synapses of both genotypes. There was no significant difference in the I/O curve basal synaptic transmission in IP_3_R2^-/-^ mice compared to their wild-type littermates IP_3_R2^+/+^ mice (**Fig. S6A**). Consistent with the above results in wild-type animals, TBS priming in SO strongly inhibited LTP in IP_3_R2^+/+^ mice (**Fig. 2C,E**). However, this effect was notably absent in slices taken from IP_3_R2^-/-^ mice while the control MML LTP remained unaffected (**Fig. 2D,E**). Importantly, in the primed pathway, the deletion of IP_3_ R2 did not affect the induction of SO LTP by the priming stimulation as measured at 30 min post-priming (**Fig. S7A**,**B**). However, similar to the effect observed in wild-type mouse hippocampal slices (**Fig. S4A**,**B**), LTP in the primed pathway underwent complete depotentiation (**Fig. S7A**,**B**). To further confirm the critical role of astrocytes in mediating trans-regional metaplasticity, we recorded field excitatory postsynaptic potentials through the astrocytic plasma membrane (AfEPSPs) through pipettes attached to single whole-cell patch-clamped astrocytes in MML. EGTA was internally delivered (calcium clamp) in a subset of recorded cells, while glycine was continuously bath-applied, as this permits LTP induction at synapses near the clamped astrocyte [19]. LTP was readily inhibited upon delivering SO priming stimulation in the absence of the calcium clamp when compared to non-primed EGTA controls (**Fig. 2E,F**). However, MML LTP post-priming was restored to control levels in the presence of the calcium clamp (**Fig. 2E,F**).

### Transregional metaplasticity triggered by either mGluR or mAChR activation

To understand the neuronal triggers for the metaplasticity effect, we began by investigating the role of conventional excitatory glutamatergic transmission using a cocktail of inhibitors for AMPARs, NMDARs, kainate receptors, and Group I/II mGluRs that was administered during the priming stimulation, followed by washout [7, 24]. To prevent residual drug effects from interfering with LTP induction in the MML, these experiments were conducted without ACSF re-circulation. Consistent with previous findings on heterodendritic metaplasticity in the SR of CA1 [5], TBS priming significantly inhibited MML LTP compared to controls, even when SO priming occurred in the presence of a cocktail of these antagonists (**Fig. S8A**). As expected, in the SO priming pathway, priming stimulation did not produce LTP when NMDARs were blocked **(Fig. S8B**). This established that the glutamatergic transmission commonly involved in triggering synaptic plasticity mechanisms was not essential for triggering the priming effect. It also confirmed that generation of LTP in the primed pathway is not necessary for the priming effect to occur in a separate subregion [5]. We then tested whether activating cholinergic pathways, specifically M1-mAChRs, was necessary and sufficient for triggering transregional metaplasticity as synaptically released acetylcholine is known to evoke Ca^2+^ elevations in astrocytes [25]. We found that pharmacological priming with bath application of an agonist of M1-mAChRs, 77-LH-28-1 oxalate (1 μM, 10 min), inhibited subsequent MML LTP 30 min later compared to DMSO controls (**Fig. 3A)**. However, contrary to our expectations and previous results [5], priming still inhibited subsequent MML LTP even in the presence of the M1 mAChR selective antagonist pirenzepine (20 μM) (**Fig. 3B**) or the non-selective mAChR antagonist atropine (10 μM) (**Fig. S9A**). Neither atropine **(Fig. S9B)** nor pirenzepine (**Fig. S9C**) affected SO LTP in the priming pathway. Thus, M1 mAChR activation is sufficient but not necessary for triggering the metaplasticity effect.

**Figure 3.**
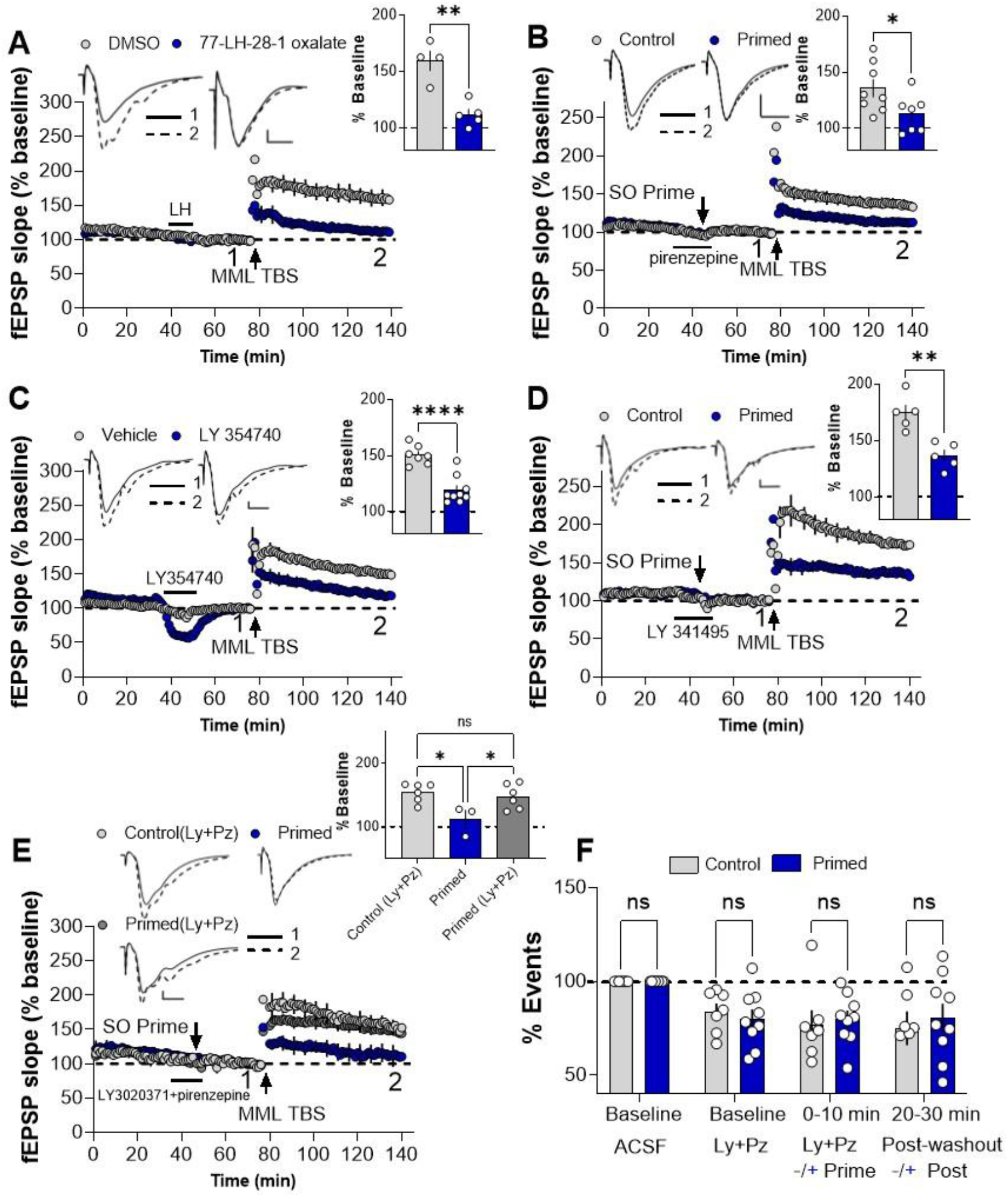
Activation of group II mGluRs and M1-mAChRs can trigger transregional metaplasticity. **(A)** Pharmacological priming with an agonist of M1-mAChRs, 77-LH-28-1 oxalate (1 μM), inhibited subsequent MML LTP (DMSO Control: 159.7 ± 8.7%, *n* = 4; LH Primed: 112 ± 4.8%, *n* = 5, *t*_*(7)*_ = 5.1, *p* = 0.0014). **(B)** However, SO TBS priming in the presence of 20 μM pirenzepine, M1-mAChR selective antagonist, did not block the transregional metaplasticity effect (Control: 135.7 ± 7.8%, *n* = 8; Primed: 113.0 ± 6.4%, *n* =7, *t*_*(13)*_ = 2.2, *p* = 0.045). **(C)** Pharmacological priming with a selective agonist of group II mGluRs, LY35470 (2 μM), inhibited subsequent MML LTP (Vehicle Control: 150.7 ± 3.2%, *n* = 7; LY Primed: 119.4 ± 4.1%, *n* = 9, *t*_*(14)*_ = 5.7, *p <* 0.0001). **(D)** However, SO TBS priming in the presence of 100 μM LY341495, a group I and II mGluR antagonist, did not block the transregional metaplasticity effect (Control: 174.5 ± 6.9%, *n* = 5; Primed: 136.4 ± 5.2%, *n* = 5, *t*_*(8)*_ = 4.4, *p* = 0.0023). The agonist-induced transient reduction in baseline is consistent with group II mGluRs acting as an autoreceptor at MPP-DG synapses [65]. **(E)** SO TBS priming in the presence of a cocktail of highly selective and potent antagonists of group II mGluRs (LY3020371, 2 μM) and M1-mAChRs (pirenzepine, 20 μM) blocked the inhibition of MML LTP seen in the absence of the drugs (one-way ANOVA *F*_*(2, 12)*_ = 5.343, *p* = 0.0219). Specifically, MML LTP post-priming with the cocktail (Cocktail Primed: 148.1 ± 8.1%, *n* = 6) was not significantly different from the non-primed control LTP (Cocktail Control: 154.5 ± 6.04%, *n* = 6, post hoc Tukey’s test, *p* = 0.82). In contrast, SO priming in the absence of the cocktail (Vehicle Primed) resulted in a significant inhibition of MML LTP (Vehicle Primed: 112.5 ± 13.8%, *n = 3*), which was significantly lower than both the drug-primed (*p* = 0.0482) and non-primed drug control groups (*p* = 0.0201). (**F**) Effects of LY3020371 (2 μM) and pirenzepine (20 μM) on Ca^2+^ calcium transients in DG MML astrocytes in response to SO priming stimulation. Two-way ANOVA indicated no significant difference between groups (*F*_*(1, 12)*_ = 0.92, *p* = 0.35), indicative of the drugs blocking the priming-induced increase in Ca^2+^ transients shown in **Fig. 2B**. Post hoc Uncorrected Fisher’s LSD tests confirmed that there were no group differences at either 0-10 min post-priming (% baseline: Control = 77.03 ± 7.6%, *n* = 7; Primed: =79.87 ± 4.5%, *n* = 9, *p* = 0.82), or 20-30 min post-priming and drug washout of the drug (Control: = 75.04 ± 9%, *n* = 7; Primed: = 80.54 ± 7.5%, *n* = 7, *t*_*(12)*_ = 1.2, *p* = 0.19). ***Inset:*** Representative waveforms are an average of 10 synaptic responses prior to LTP induction (**1**) and at the conclusion of the experiment (**2**). Bar graphs summarize LTP across individual slices. Scale bars: 0.5 mV, 5 ms (**A**), 1 mV, 5 ms (**B**), 0.5 mV, 10 ms (**C-E**). Arrows indicate the timing of SO priming and MML LTP induction. All data presented as mean ± SEM; *ns, p* > 0.05; ∗, *p* < 0.05; ∗∗, *p* < 0.001, ∗∗∗∗, *p* < 0.0001.

We then asked what other receptor system could be involved in the priming effect? Previous studies have shown that the activation of group II mGluRs can modulate both synaptic plasticity and astrocyte Ca^2+^ activity [26-28]. We thus tested whether the group II mGluR agonist LY35470 alone could mimic a priming effect on MML LTP. Indeed, pharmacological priming with LY35470 (2 μM) inhibited MML LTP to a similar extent as electrical priming in SO (Fig. 3C). However, electrical priming in SO still inhibited subsequent MML LTP compared to controls when given in the presence of the mGluR antagonist LY341495 used at a concentration (100 μM) that blocks all mGluR subtypes (Fig. 3D). Since neither Group II mGluR antagonist (MCPG and LY341495) blocked the electrical priming effect, these results indicate that Group II mGluR stimulation is also sufficient but not necessary for triggering the metaplasticity.

The parallel findings for the agonists and inhibitors for both Group II mGluRs and M1-mAChRs suggested that either pathway can initiate the priming effect and that electrical priming in SO activates both pathways simultaneously. To test this hypothesis, we blocked both signaling pathways during electrical priming. The co-application of either LY341495 (100 μM) and atropine (10 μM) (**Fig. S10A**), or the more highly selective and potent blockers of group II mGluRs, LY3020371 (20 μM), and M1-mAChRs, pirenzepine (20 μM) (**Fig. 3E**), prevented the inhibition of MML LTP. This co-application did not affect the homosynaptic LTP in the SO priming pathway (**Fig. S10B**,**C**).

To confirm that the neurotransmitters glutamate and acetylcholine activate astrocytes, we once again performed Ca^2+^ imaging in MML astrocytes in response to SO electrical priming, this time in the presence of the cocktail of LY3020371 (20 μM) and pirenzepine (20 μM). Notably, there was no difference in the frequency of astrocytic Ca^2+^ events immediately following priming (**SV 2**) or 20 min post-drug washout (**SV2**) compared to non-primed controls under these conditions (**Fig. 3F**), in contrast to the increase reported above in the absence of the antagonists (**Fig. 2A**). We also investigated the role of purinergic signaling, specifically through P2Y1 receptors (P2Y1Rs), in the transregional metaplasticity. Neuronal ATP activates astrocytic P2Y1Rs and astrocytic ATP release through connexin hemichannels, resulting in astrocytic Ca^2+^ signals, highlighting a potential role for purinergic signaling [29-32]. Connexin-43 is already known to be crucial for heterodendritic metaplasticity in SR [8]. Here, we found that applying the P2Y1R antagonist MRS 2719 (10 μM) only during SO priming did not block the subsequent inhibition of MML LTP (**Fig. S11A**). However, the bath application of MRS 2719 throughout the entire recording period did block the SO priming-mediated inhibition of MML LTP (**Fig. S11B**). This suggests that ongoing release of ATP post-priming plays a role in maintaining the metaplasticity effect across time, consistent with the ongoing elevation in the frequency of astrocyte calcium events described above.

### Role of astrocytic TNF in transregional metaplasticity

Having established an essential role of astrocytic Ca^2+^ signaling, triggerable by activation of either mGluRs or mAChRs, in transregional metaplasticity, we next investigated which specific gliotransmitter could be mediating this long-distance effect. Previously, we reported that a 10 min bath application of 1.18 nM TNF protein significantly inhibited LTP induced 30 min later in the SR of CA1 [33]. Similarly, we found that priming with the same TNF regimen significantly inhibited subsequent LTP in the MML in rat hippocampal slices (**Fig. S12A**).

The next step was to identify the cellular source of this TNF, as it is known to be produced in the central nervous system by various cell types, including neurons, microglia, and astrocytes[34]. To pinpoint the specific cell type responsible for TNF release in our context, we conducted experiments using transgenic mice with TNF knocked out in either microglia or astrocytes. The experimental setup was slightly different from most previous studies (but see [5], as the timing between priming and MML LTP recording was reduced to 15 min. First, we performed SO electrical priming in Cx3cr1Cre^+/−^; TNF^flox/flox^ mice, where TNF is specifically knocked out in microglia [15]. In these mice, SO priming still impaired subsequent MML LTP compared to non-primed controls (**Fig. 4A**). This finding strongly suggests that microglia are not the primary source of TNF release. However, when a similar set of experiments was undertaken in GFAP Cre^+/-^; TNF^flox/flox^ mice, where TNF expression was ablated specifically in astrocytes [15], the SO priming effects on MML LTP were completely blocked. In contrast, their littermate GFAP Cre^-/-^; TNF^flox/flox^ mice still exhibited a significant reduction in subsequent MML LTP (**Fig. 4B,C**). To confirm these findings, we used a conditional astrocyte-specific TNF knockout in older mice (4-6 months old) using tamoxifen-inducible Aldh1l1−Cre/ERT2^+/-^ mice crossed with TNF^flox/flox^ mice (**Fig. 4D**). The successful deletion of TNF in astrocytes was qualitatively confirmed by visualizing the TdTomato reporter in astrocytes (**Fig. S13**). Consistent with the prior result, the metaplastic inhibition of MML LTP was absent in the Aldh1l1−Cre/ERT2Cre^+/−^; TNF^flox/flox^ mice, whereas the Aldh1l1−Cre/ERT2^−/−^; TNF^flox/flox^ mice (TAM-injected control littermates) still showed a significant priming effect (**Fig. 4E,F**).

**Figure 4.**
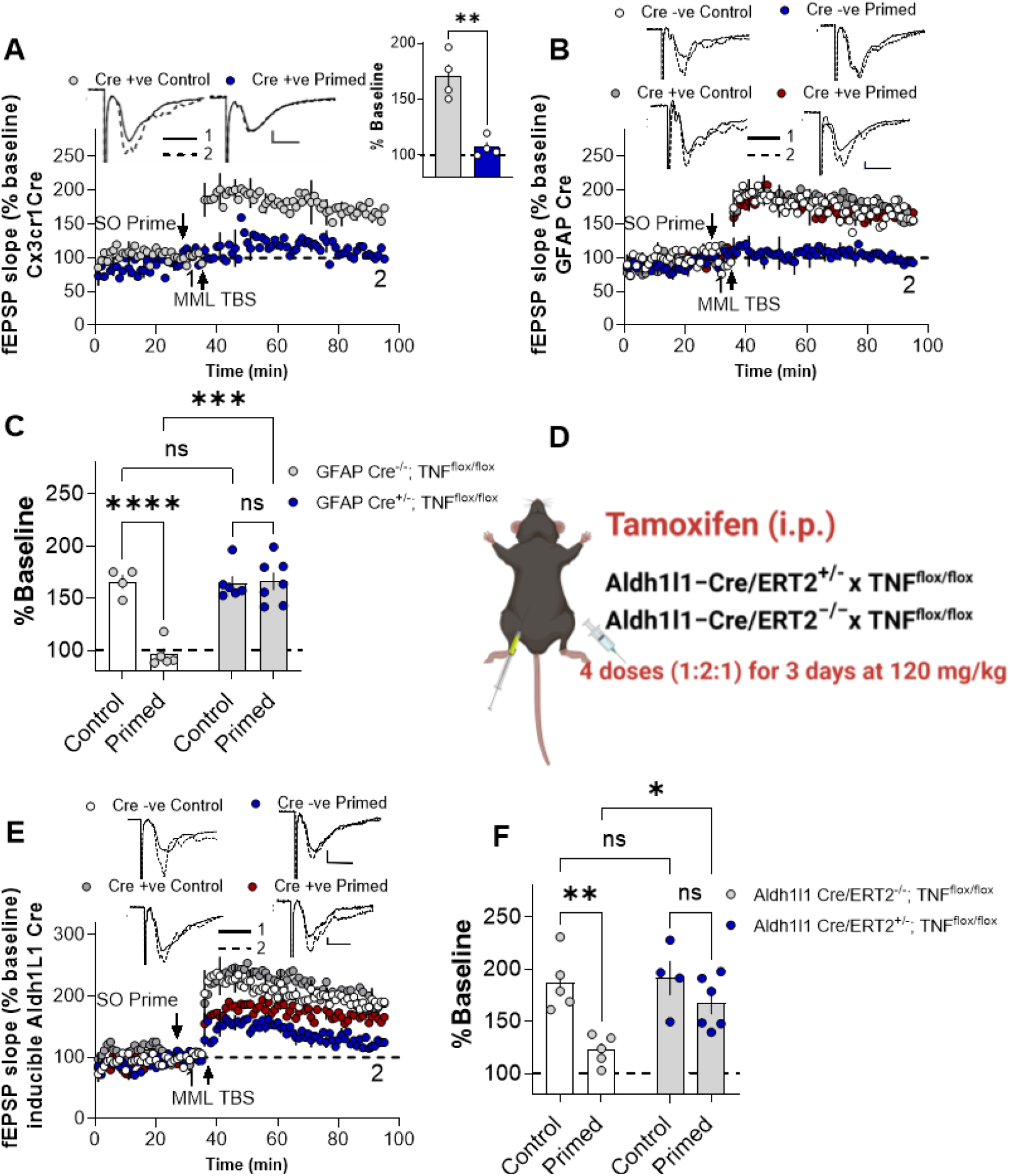
Astrocytes are the cellular source of TNF mediating transregional metaplasticity. **(A)** Microglia are not the cellular source of TNF in mediating trans-regional metaplasticity. SO electrical priming in Cx3-Cr1Cre^+/-^ TNF^flox/flox^ mice resulted in a significant reduction in subsequent MML LTP compared to non-primed controls (Control non-primed: 170.6 ± 10.3%, *n* = 4; Primed: 107.2 ± 4.4%, *n* = 4, *t*_*(6)*_ = 5.7, *p* = 0.0013). **(B-C)** Astrocytes are the cellular source of TNF in mediating transregional metaplasticity. Two-way ANOVA revealed a significant priming x genotype interaction effect (*F*_*(1, 7)*_ = 23.76, *p* = 0.0018). The SO priming effect on MML LTP was completely prevented in GFAP Cre^+/-^ TNF^flox/flox^ (Control non-primed: 164.3 ± 6.7%, *n* = 6; Primed: 166.3 ± 8.1%, *n* =7, post-hoc Uncorrected Fisher’s LSD test, *p* = 0.83) compared to GFAP Cre^-/-^ TNF^flox/flox^ that showed a significant reduction in subsequent MML LTP (Control: 165.8 ± 6.4%, *n* = 4; Primed: 96.1± 5.6%, *n* = 5, *p* < 0.0001). **(D)** Schematic showing the tamoxifen administration for the Aldh1l1−Cre/ERT2^+/−^or Aldh1l1−Cre/ERT2^−/−^ crossed with TNF^flox/flox^ transgenic mice. **(E-F)** Tamoxifen-induced knock-out of TNF in astrocytes also restricted the subsequent inhibition of MML LTP upon SO priming. Two-way ANOVA indicated a significant priming x genotype interaction effect (*F*_*(1, 7)*_ = 3.401, *p* = 0.1077). The SO electrical priming effect on subsequent MML LTP was significantly blocked in the Aldh1l1-Cre/ERT2 Cre^+/-^ TNF^flox/flox^ (Control non-primed: 191.5 ± 16.1%, *n* = 4; Primed: 167.3± 10.2%, *n* = 6, post hoc Uncorrected Fisher’s LSD test, *p* = 0.1499) while the Aldh1l1-Cre/ERT2 Cre^-/-^; TNF^flox/flox^ mice showed a significant reduction in MML LTP after priming (Control non-primed: 186.9 ± 12.3%, *n* = 5; Primed: 122.6 ± 6.36%, *n* = 5, *p* = 0.0010). ***Inset:*** Representative waveforms are an average of 10 synaptic responses prior to LTP induction (**1**) and at the conclusion of the experiment (**2**). Bar graphs summarize LTP across individual slices. Scale bars: 0.5 mV, 5 ms (**B-C**), 1 mV, 5 ms (**E, top panel**), 0.5 mV, 5 ms (**E, bottom panel**). Arrows indicate the timing of SO priming and MML LTP induction. All data presented as mean ± SEM; ns, *p* > 0.05; ∗, *p* < 0.05; ∗∗, *p* < 0.01, ∗∗∗, *p* < 0.001, ∗∗∗∗, *p* < 0.0001.

We then asked whether TNF was acting specifically at TNF type 1 receptors (TNFR1), as it does for the priming effect in CA1 SR [6]. We first performed TNF protein priming in mice lacking TNFR1 (TNFR1^-/-^ mice), alongside their wild-type littermates (TNFR1^+/+^ mice). Pharmacological priming with 1.18 nM TNF significantly inhibited MML LTP in TNFR1^+/+^ mice (**Fig. 5A,C**), but not in the TNFR1^-/-^ mice (**Fig. 5B,C**). To confirm the necessity of TNFR1 activation, we undertook SO electrical priming in mouse hippocampal slices from TNFR1^-/-^ mice and their TNFR1^+/+^ littermates. SO TBS priming readily inhibited subsequent MML LTP in TNFR1^+/+^ mice (**Fig. 5D,F**), but this effect was absent in the TNFR1^-/-^ mice (**Fig. 5E,F**).

**Figure 5.**
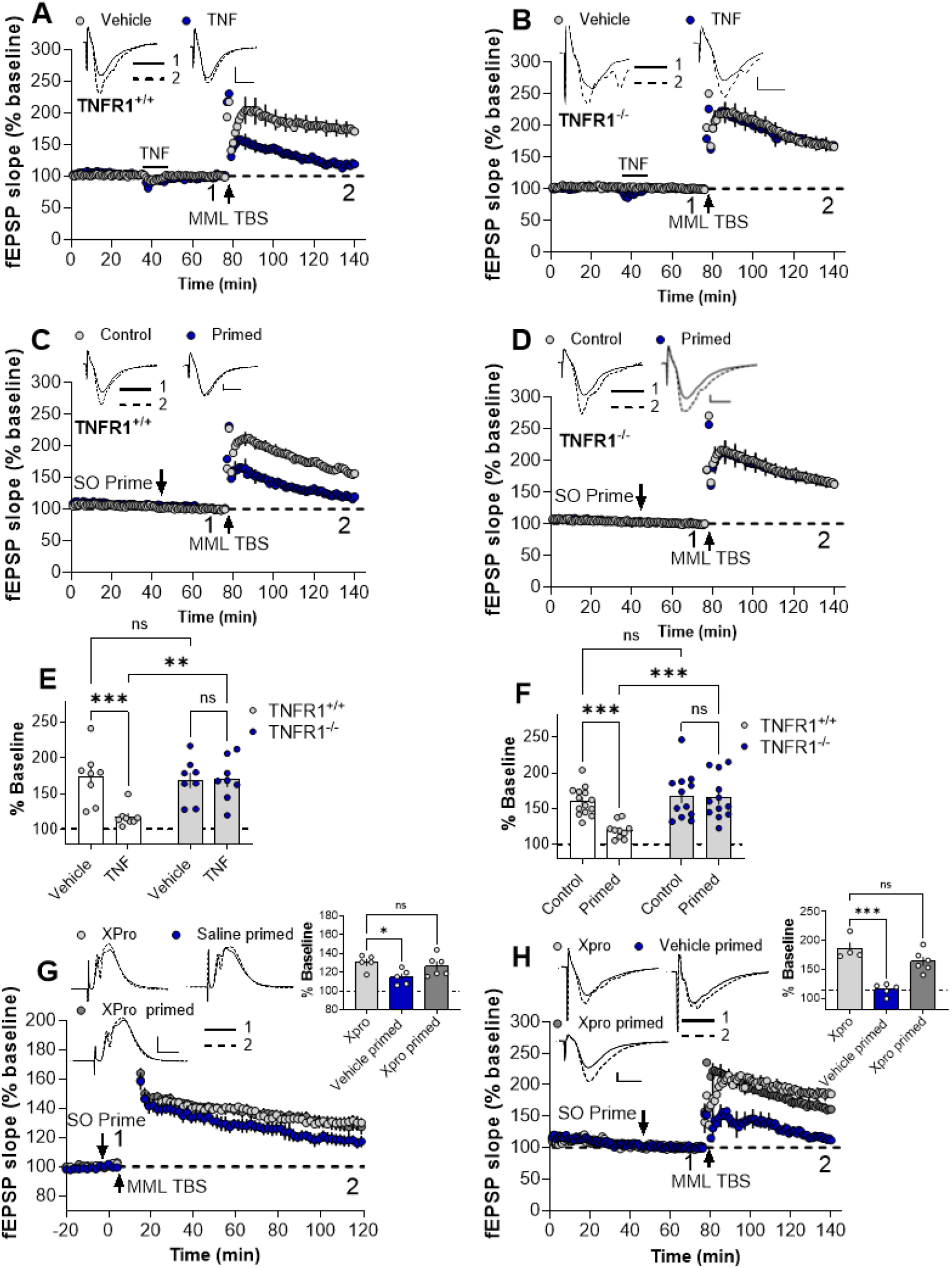
TNF acts via TNFR1 transregional metaplasticity. **(A-B,E)** TNF priming requires activation of TNFR1. Two-way ANOVA indicated a significant TNF priming x genotype interaction effect (*F*_*(1, 14)*_ = 8.9, *p* = 0.0096). Pharmacological priming using 1.18 nM TNF significantly inhibited MML LTP in **(A**,**E)** TNFR1^+/+^ mice (Vehicle Control: 173.9 ± 13%, *n* = 8; TNF Primed: 117.9 ± 4.8%, *n* = 8, post hoc Uncorrected Fisher’s LSD test, *p* = 0.0006), but not in the (**B**,**E**) TNFR1^-/-^ mice (Vehicle Control: 169 ± 10.6%, *n* = 8; TNF Primed: 170.3 ± 10.7%, *n* = 8, *p* = 0.93). **(C-D**,**F)** SO TBS priming-mediated metaplasticity requires activation of TNFR1. Two-way ANOVA indicated a significant electrical priming x genotype interaction effect (*F* _*(1, 20)*_ = 7.6, *p*=0.0122). Thus, the priming in SO readily inhibited subsequent MML LTP in **(C)** TNFR1^+/+^ mice (Control non primed: 160.4 ± 5.3 %, *n* =14; Primed: 120.1 ± 3.6 %, *n* =10, post hoc Uncorrected Fisher’s LSD test, *p =* 0.0004), but not in the **(D)** TNFR1^-/-^ mice (Control non-primed: 167.0 ± 9%, *n* = 13; Primed: 166.1± 8.2%, *n =* 13, *p* = 0.80). **(G-H)** XPro1595, a potent and selective inhibitor of soluble TNF that binds to TNFR1, blocked the SO priming effects on subsequent MML LTP. One-way ANOVAs indicated a significant main effect of group, both **(G)** *in vivo* (*F*_*(2, 13)*_ = 3.962, *p* = 0.045) and **(H)** *in vitro* (*F*_*(2, 12)*_ = 21.7, *p* = 0.0001). Thus, XPro1595 (10 mg/kg, s.c.) administered 12-16 hours earlier prevented SO priming-mediated inhibition of MML LTP *in vivo* in anesthetized rats (Control non-primed: 130.7 ± 3.5%, *n* = *5*; Primed: 126.8 ± 4.4%, *n =* 6, post hoc Tukey’s test, *p* = 0.77). However, SO priming in the absence of XPro1595 inhibited MML LTP (Primed: 115.0 ± 3.6%, *n = 5, p =* 0.046). (**H**) Similar results were obtained *in vitro* in mouse hippocampal slices (Control non-primed: 186 ± 10.2%, *n* = 4; Primed: 164 ± 7.5%, *n* = 6, post hoc Tukey’s test, *p* = 0.14). However, SO priming in the absence of XPro1595 inhibited MML LTP (Primed: 115.5 ± 4.5%, *n = 5*) when compared to LTP measured in the presence of XPro1595 priming (*p* = 0.0012) and in the absence of priming (*p* = 0.0001). ***Inset:*** Representative waveforms are an average of 10 synaptic responses prior to LTP induction (**1**) and at the conclusion of the experiment (**2**). Bar graphs summarize LTP across individual slices. Scale bars: 0.5 mV, 5 ms (**A-B, D, H**), 1 mV, 5 ms (**C**), 5 mV, 5 ms (**G**). Arrows indicate the timing of SO priming and MML LTP induction. All data presented as mean ± SEM; ns, *p* > 0.05; ∗, *p* < 0.05; ∗∗, *p* < 0.001, ∗∗∗, *p* < 0.001.

In the SO priming pathway, TBS priming resulted in substantial potentiation in both TNFR1^+/+^ mice and TNFR1^-/-^ mice, as measured at 30 minutes post-priming (**Fig. S14A**,**B**). However, as reported previously, the SO LTP in the TNFR1^+/+^ mice underwent complete depotentiation, such that it was not significantly different from the non-primed control slices (**Fig. S14A**). Interestingly, unlike in the TNFR1^+/+^ mice, the depotentiation of SO LTP was not observed in TNFR1^-/-^ mice (**Fig. S14B**). The reason for this difference remains unclear.

Finally, to reinforce TNF’s role in acting on TNFR1s, we investigated whether blocking the binding of soluble TNF to TNFR1 using XPro1595 [20] would mitigate the SO priming effect on MML LTP. We observed that injection of XPro1595 on the day before experimentation prevented the priming effect on subsequent MML LTP in both anesthetized rats (**Fig. 5G**) and mouse hippocampal slices (**Fig. 5H**), restoring LTP almost to the level observed in vehicle-treated controls (**Fig. 5G,H)**. XPro1595 also partially blocked the depotentiation effect seen in mouse SO following MML tetanization (**Fig. S15**).

### Activation of GluN2B-containing NMDARs and transregional metaplasticity

It is known that TNF can trigger Ca^2+^-dependent glutamate release from astrocytes, which in turn modulates excitatory synaptic activity in the hippocampus, including at granule cell extrasynaptic NMDARs [10-12, 35]. Accordingly, we investigated the requirement for GluN2B-containing NMDAR activation in the transregional metaplasticity. Priming experiments were conducted under two distinct conditions: 1) with the GluN2B antagonist ifenprodil (0.8 μM) bath-applied throughout the entire experiment, or 2) with ifenprodil bath-applied beginning 5 minutes post-priming and then for the rest of the experiment.

In the continuous presence of 0.8 μM ifenprodil, the priming-mediated inhibition of MML LTP was blocked (**Fig. 6A**), while the non-primed control LTP in MML remained unaffected by the ifenprodil. In the SO priming pathway, TBS induced substantial homosynaptic LTP that was unaffected by ifenprodil (**Fig. S16A**). However, as shown in **Fig. S8A**, NMDAR activation during the priming stimulation itself is not required for the transregional metaplasticity. Therefore, any potential NMDAR involvement was likely to occur during the interval between priming and LTP induction. To test this hypothesis, we washed in ifenprodil 5 minutes post-priming. Even under these conditions, ifenprodil still blocked the inhibition of MML LTP (**Fig. 6B**), while SO LTP again remained unaffected (**Fig. S16B**).

**Figure 6.**
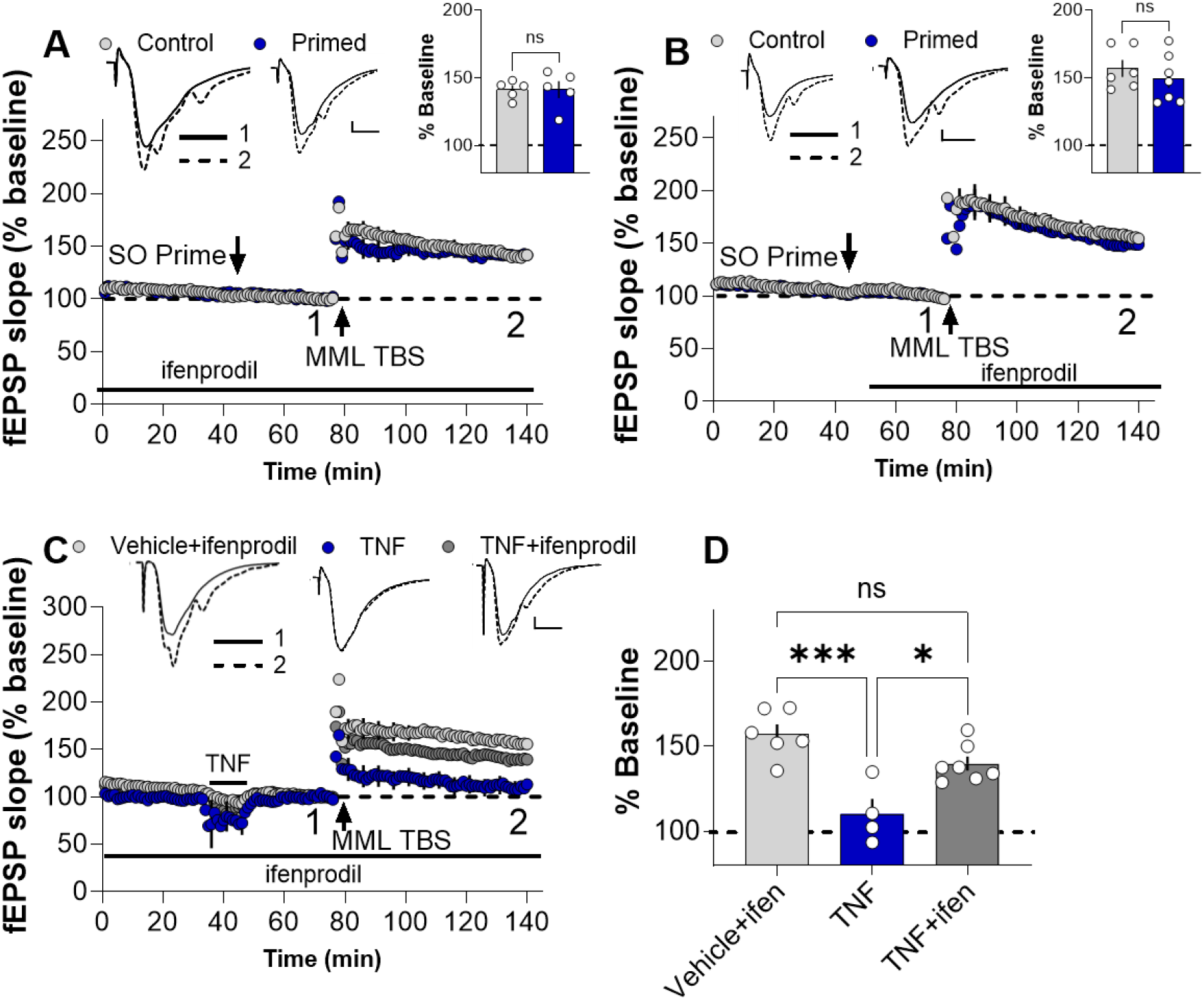
Transregional metaplasticity requires activation of GluN2B-containing NMDA receptors. **(A-B)** Priming SO in the presence of 0.8 μM ifenprodil blocked the inhibition of MML LTP. (**A**) Bath application of ifenprodil throughout the experiment blocked the priming effect on MML LTP (Control non-primed: 141.3 ± 3%, *n* = 5; Primed: 141.4 ± 6.2%, *n* = 5, *t*_*(8)*_ = 0.022, *p* = 0.98). **(B)** Bath-applied ifenprodil, just after SO priming, also blocked the inhibition of MML LTP (Control non-primed: 156.9 ± 6.3%, *n* = 6; Primed: 149.2 ± 6.7%, *n* = 7; *t*_*(11)*_ = 0.83, *p* = 0.43). **(C-D)** Ifenprodil blocked TNF priming-mediated inhibition of MML-LTP (one-way ANOVA, *F* _*(2, 14)*_ = 13.96, *p* = 0.0005). Thus, there was no difference in MML LTP between the Vehicle+ ifenprodil group (157.3 ± 5.7%, *n* = 6) and the TNF Primed+ ifenprodil group (139.8 ± 4.2%, *n* = 7, post-hoc Tukey’s, *p* = 0.0918). In the absence of ifenprodil, TNF priming (1.18 nM) still inhibited subsequent MML LTP (TNF Primed: 110.3 ± 8.9%, *n* = 4) compared with MML LTP measured in the presence of ifenprodil (*p* = 0.0110) and in the absence of priming (*p* = 0.0003). **(D)** Bar graph summarizing ANOVA results, TNF priming effects in the presence and absence of ifenprodil. ***Inset:*** Representative waveforms are an average of 10 synaptic responses prior to LTP induction (**1)** and at the conclusion of the experiment (**2**). Inset bar graphs summarize LTP across individual slices. Scale bars: 0.5 mV, 5 ms (**A-C**). Arrows indicate the timing of SO priming and MML LTP induction. All data presented as mean ± SEM; ns, *p* > 0.05; ∗, *p* < 0.05; ∗∗∗, *p* < 0.001

To investigate whether TNF acting on TNFR1 is upstream of glutamate acting on GluN2BRs, we undertook TNF priming in the ACSF containing 0.8 μM ifenprodil. As for electrical priming, ifenprodil blocked the TNF priming effect (**Fig. 6C**), while in the absence of ifenprodil, TNF priming inhibited subsequent MML LTP (**Fig. 6C**). These results indicate that GluN2B-containing NMDARs mediating the inhibition of LTP are downstream of TNF signaling in this transregional metaplasticity pathway.

## Discussion

This study has revealed a novel transregional metaplasticity effect within the hippocampus, whereby neural activity originating in the outer dendritic layer of CA1 can profoundly influence synaptic plasticity at MML synapses in the DG. This effect, spanning hundreds of microns away from the site of simulation and across the hippocampal fissure, challenges the traditional view of information flow within the trisynaptic circuit by highlighting a previously unrecognized level of inter-regional coordination. The observed inhibition of MML LTP, induced by both HFS and TBS priming in CA1, was robust across species, sexes, and tissue preparations, suggesting a fundamental physiological mechanism. Critically, this effect was not solely dependent on direct neuronal connections but required an elaborate signaling cascade starting with activation of Group II mGluR and/or M1-mAChRs. These receptors are likely the ones expressed by astrocytes, since their activation triggered an IP_3_R2-dependent astrocytic Ca^2+^ signaling cascade, ultimately leading to the release of TNFR1, and then glutamate acting on GluN2B-containing NMDARs. These NMDARs are likely extrasynaptically located, as their activation is known to promote LTD and impair LTP [36], a pattern consistent with our priming effects [5]. Our findings significantly expand on the understanding of astrocytic contributions to higher-order function beyond their established roles in local modulation of synaptic plasticity.

The most striking aspect of this transregional metaplasticity is its ability to operate across the hippocampal fissure, a region traditionally considered a physical barrier with very limited direct neuronal innervations between CA1 and DG. However, astrocytes are known to communicate across the fissure [9]. Astrocytes inhabit non-overlapping but connected territories, in which each astrocyte may contact up to 140,000 synapses [37]. This structural arrangement allows astrocytes to uniquely sense nearby cellular activity and respond by modulating the induction of either homosynaptic or heterosynaptic LTP and LTD, depending on the type of gliotransmitter released [38, 39]. The observation that priming stimulation in CA1 SO triggers Ca^2+^ events in the DG MML astrocytes, across the hippocampal fissure, confirms the unique capacity of astrocytic networks to connect anatomically distinct sub-regions of the hippocampus. The pivotal role of IP_3_R2-mediated Ca^2+^ release from astrocytic intracellular stores underscores the importance of astrocytic signaling for initiating these transregional effects. In complementary work, previous studies reported significant impairments both in long-term memory consolidation and LTD in the CA1 SR of the hippocampus of IP_3_R_2_^-/-^ mice [40]. These findings align with our previous report demonstrating that SO priming not only inhibited subsequent LTP but also enhanced LTD in the CA1 SR [5]. Together, these findings shift the perspective of astrocytes from being viewed merely as local regulatory and support cells to recognizing them as active participants in orchestrating large-scale communication and information processing across neuronal networks.

The necessity of either acetylcholine acting on M1-mAChRs or glutamate on group II mGluRs to trigger the astrocytic Ca^2+^ signaling could suggest that such neuronal activity in CA1 directly targets nearby astrocytes [41, 42]. Astrocytes express Group II mGluRs and contribute to Ca^2+^ signaling [27] from intracellular stores via the G_β_γ subunit, directly activating phospholipase C or through interactions with IP_3_ receptors [43-45]. Likewise, astrocytes express m1-AChRs and respond to both acetylcholine [46] and the activation of cholinergic fibers present in SO [25, 47]. This response to M1-mAChR activation occurs via Gq signaling by increasing Ca^2+^ activity in an IP_3_R2-dependent manner and releasing gliotransmitters that regulate neuronal function [48, 49]. The observation that purinergic signaling via P2Y1Rs, while essential, may not be the initial trigger but rather facilitates the spatial spread and temporal persistence of Ca^2+^ signaling in astrocytes [30], further emphasizes the complex nature of neuron-astrocyte communication in the regulation of synaptic plasticity. Previous reports suggest that activation of P2Y1Rs might be critical for astrocytic Ca^2+^ signaling, which in turn facilitates the release of glutamate from astrocytes, a process found to be blocked by P2Y1 receptor antagonists [35]. This aligns well with our finding that prolonged but not transient application of P2Y1R antagonists blocked the priming effects on MML LTP, suggesting their involvement in later stages of the signaling cascade.

The release of TNF from astrocytes and its action on TNFR1s represents a gliotransmitter-mediated communication pathway that contributes to the metaplasticity signaling mechanisms. Interestingly, we have shown using pharmacological priming that TNF likely acts in an autocrine fashion back on astrocytes as TNF priming-mediated inhibition of LTP in CA1 SR was blocked by buffering Ca^2+^ in CA1 SR astrocytes [7]. These observations indicate that TNF is not the final signal released onto neurons. Rather, glutamate acting on GluN2B-containing NMDARS appears to be the final signal causing the downregulation of MML LTP since TNF priming was blocked by ifenprodil (**Fig. 6C**). Already, roles of GluN2B-containing NMDARs in mediating metaplasticity have been reported in the hippocampus. For example, prior activity modifies synaptic thresholds with quiescent synapses showing greater structural growth and LTP, as triggered by activity-dependent remodeling of GluN2B-subunit expression [50]. Further, astrocytes release glutamate that targets neuronal receptors in dendrites or axonal terminals for neuromodulatory actions [11]. In CA1, astrocytic glutamate acts on extrasynaptic GluN2B-containing NMDARs located on the dendrites of CA1 pyramidal cells. Thus, astrocyte-dependent glutamate release, by acting on extrasynaptic GluN2B-sensitive receptors, critically contributes to the metaplastic modulation of LTP, that occurs in a spatially precise manner across different hippocampal subregions [51]. The hippocampus comprises a largely unidirectional trisynaptic circuit, which is crucial for the formation and processing of episodic and spatial memories [52]. The DG and CA1 play distinct yet interconnected roles in memory processing. The DG is considered the first relay in the trisynaptic circuit, and is thought to be involved in pattern separation, ensuring that similar inputs are encoded as distinct memories [53, 54]. CA1, receiving highly processed information from CA3, is considered critical for pattern completion and memory retrieval [55]. The existence of transregional metaplasticity suggests a mechanism by which activity in CA1 can dynamically regulate the LTP threshold in the DG. This “top-down” modulation from CA1 to DG could serve as a higher-order regulatory mechanism for memory encoding [56, 57]. This higher-order control could serve to stabilize the network, whereby successful memory retrieval in CA1 signals the DG to become temporarily less plastic to prevent interference from new, similar inputs or, conversely, ensuring only truly novel or stronger signals trigger new encoding, thereby maintaining global network homeostasis [58, 59]. Importantly, this mechanism is a compelling example of highly selective long-distance information transfer, as the priming signal impacts SR synapses but avoids other synapses in CA1 [6], while also targeting the crucial MPP inputs to the DG MML. Thus, this regulatory system is not simply a non-selective broadcast but rather targets specific downstream synaptic populations [60]. This capacity for constant crosstalk and dynamic regulation highlights an integrative mechanism such that memory traces are not formed serially but refined through inter-regional communication. Crucially, this dynamic regulation involves a neuron-glia circuit wherein astrocytes mediate the long-distance signal as they discriminate and integrate activity across different pathways [28, 41, 61].

Dysregulation of synaptic plasticity is a hallmark of numerous neurological and psychiatric disorders, including Alzheimer’s disease (AD), epilepsy, and schizophrenia. We have previously shown that the heterodendritic metaplasticity mechanism is aberrantly engaged in the APP/PS1 mouse model of AD in a TNF-dependent manner, such that there is impaired LTP in CA1 SR even in the absence of experimentally provided priming stimulation, an effect that can be rescued by a TNF antibody [33]. Our understanding of transregional metaplasticity, mediated by astrocytes and TNF, opens new avenues for understanding disease pathogenesis and developing therapeutic interventions. In AD, for instance, neuroinflammation and astrogliosis are prominent features, and elevated TNF levels are frequently observed both in mice and human patients [33, 62]. Aberrant TNF signaling, primarily through the sustained activation of its neurotoxic TNFR1 receptor, is a core mechanism of synaptic dysfunction and cognitive decline across multiple neurological disorders, including AD, multiple sclerosis, and acute conditions like traumatic brain injury [20, 63, 64]. In these conditions, chronically elevated levels of TNF act as a strong driver of neuroinflammation, leading to gliosis and neurotoxic changes that negatively impact neuronal function. This dysregulated signaling directly disrupts synaptic plasticity and metaplasticity, primarily by suppressing LTP. As a result, there is impaired memory formation and associated cognitive deficits.

## Materials and Methods

### Animals and slice preparation

All procedures were approved by the University of Otago Animal Ethics Committee and complied with New Zealand Animal Welfare Legislation. Male Sprague-Dawley rats (6-8 weeks old, obtained from breeding facility at University of Otago) were deeply anesthetized with ketamine (100 mg/kg, i.p.) and decapitated. Brains were rapidly dissected and submerged in ice-cold sucrose dissection solution (in mM: 210 sucrose, 26 NaHCO_3_, 2.5 KCl, 1.25 NaH_2_PO_4_, 0.5 CaCl_2_, 3 MgCl_2_, 20 D-glucose, saturated with 95% O_2_/5% CO_2_). Hippocampi were isolated, and CA3 was removed before transverse hippocampal slices (400 μm) were prepared using a Leica VT 1000 S vibratome. Slices were incubated at interface with artificial cerebrospinal fluid (ACSF, in mM: 124 NaCl, 3.2 KCl, 1.25 NaH_2_PO_4_, 26 NaHCO_3_, 2.5 CaCl_2_, 1.3 MgCl_2_, 10 D-glucose) at 32°C for 30 min, followed by room temperature until use. Post-incubation slices were submerged in a humidified recording chamber perfused with ACSF at 2.5 ml/min and a temperature of 32.5°C ± 0.2 °C. Unless otherwise indicated, ACSF was recycled through the course of experimental recording.

For mouse hippocampal slice preparation, adult wild-type C57BL/6J or transgenic mice (males and females, 2-7 mo) were anesthetized with isoflurane and decapitated. Horizontal brain sections (350-400 μm) containing the hippocampus and cortex were prepared. Mouse slices were treated similarly to rat slices, except that CA3 was left intact.

Dr. Ju Chen at the University of California, San Diego, kindly provided the *IP*_*3*_*R2*^*-/-*^ (knockout, KO) mice. The *IP*_*3*_*R2*^*-/-*^ mice were bred with female C57BL/6 mice at the University of Otago animal facility to obtain IP_*3*_*R2*^*+/-*^ (heterozygous) mice. The *IP*_*3*_*R2*^*+/-*^ mice were crossed to obtain *IP*_*3*_*R2*^*+/+*^ and *IP*_*3*_*R2*^*-/-*^ littermate mice. *TNFR1*^*-/-*^ (KO) male mice were obtained from the Jackson Laboratory (Strain#:002818) and were bred with female C57BL/6J mice present at the University of Otago animal facility to produce *TNFR1*^*+/-*^ mice (heterozygous). *TNFR1*^*+/-*^ mice were crossed to produce *TNFR1*^*+/+*^ (WT), *TNFR1*^*+/-*^, and *TNFR1*^*-/-*^ littermate mice. Both males and females were used, and the experimenter was blind to the animals’ genotype, which was only known upon completing the data collection and analysis.

For calcium imaging experiments, 6-8-week-old mice expressing *Aldh1L1-cre/ERT2* ^*+/-*^ *GCaMP6f*^*+/-*^ were administered tamoxifen (TAM, 80-120 μg/g in corn oil, i.p.) in four injections over three days or five injections over five days to achieve GCaMP6f expression in astrocytes using previously described methods [13, 14]. For all the transgenic strains, the tissue samples were outsourced to Transnetyx (TN, USA) for genotyping, a service with high accuracy. For glia-specific TNF^-/-^mice, floxed TNF mice were crossed with GFAP-Cre (astrocytic) mice or with CX3CR1-Cre (microglial) MW126Gsat mice. GFAP-Cre ^+/-^ or CX3CR1-Cre^+/-^expressing mice were compared with GFAP-Cre^-/-^ or CX3CR1-Cre^-/-^ non-expressing littermates. Floxed TNF and GFAP-Cre mice were on a C57BL/6 background; Cx3Cr1-Cre mice were on a mixed FVB/B6/129/Swiss/CD1 background. In addition, for the inducible models, a similar TAM administration described above was followed to generate TNF^flox/flox^; Aldh1L1Cre^+/−^, which was specific to astrocytes [15]. For these experiments, all animal procedures were performed at McGill University in accordance with the guidelines of the Canadian Council on Animal Care and the Animal Care Committee of the Montreal General Hospital Facility.

### Slice field electrophysiology

Field excitatory postsynaptic potentials (fEPSPs) were recorded to assess synaptic efficacy. Stimulation of afferents from CA2/CA3 and other regions running in CA1 SO as well as stimulation of medial perforant path (MPP) afferents running in the DG middle molecular layer (MML) was performed using custom-built constant current programmable stimulators (diphasic pulses, 0.1 ms half-wave duration) and 50 μm diameter Teflon-insulated tungsten monopolar electrodes. Recording electrodes (glass capillary micropipettes from AM systems, pulled to 2.0-3.5 MΩ, filled with ACSF) were placed ∼400 μm from the stimulating electrode in both strata (**Fig. S1A**). Signals were amplified (Grass® P511 AC amplifiers, 0.3 Hz-3 kHz half-amplitude filters) and digitized (National Instruments BNC-2110).

Slices were included if afferent pathway stimulation elicited fEPSPs ≥ 1 mV (SO) or ≥ 2 mV (MML) at 100 μA. To confirm the stimulation of MPP inputs, a paired-pulse stimulation test was conducted at a 50-ms inter-pulse interval in the DG both *in vitro* (**Fig. S1B)** and *in vivo* **(Fig. S1C)**. It is well established that paired-pulse stimulation typically results in paired-pulse depression (PPD) specifically when the medial perforant path is stimulated [16, 17]. Accordingly, when two pulses delivered 50 ms apart generated PPD, this was attributed to the electrode being correctly placed amongst the MPP inputs in the MML. A paired-pulse test was performed across all experimental recordings from both rat and mouse hippocampal slices using a stimulus intensity that elicited fEPSPs that were not contaminated with population spikes.

Baseline stimulation (set at 40-50% of the fEPSP slope obtained at 200 μA) was delivered every 30 s, alternating between SO and MML. Control experiments included 75 minutes of baseline recording, LTP induction, and 60-90 minutes of post-tetanization recording. Slices with baseline response changes >10% were excluded. SO priming protocols included: 1) 2 sets of 3 trains of high-frequency stimulation (HFS; 100 Hz, 1 s, 20 s intervals between trains in a set, 5 min interval between sets) or 2) 2 trains of theta-burst stimulation (TBS; 10 bursts of 5 pulses at 100 Hz, 100 μs pulse duration, 200 ms interburst intervals and 20 s intertrain interval). LTP induction in DG MML used 4 trains of TBS (10 bursts of 10 pulses at 100 Hz, 200 μs pulse duration, 200 ms interburst intervals, 30 s intertrain intervals) in the presence of 0.2 μM gabazine to reduce GABA_A_ receptor-mediated inhibition in DG. For certain transgenic mice, input-output (I/O) curves were generated at MML synapses by delivering a series of stimulations to the MPP afferent fibers using current increments (μA: 10, 20, 50, 100, 150, 200) and recording the corresponding fEPSP responses.

### *In vivo* electrophysiology

Male Sprague-Dawley rats weighing between 400-700 g were anesthetized with urethane (1.5 g/kg, i.p.) and surgically prepared with indwelling electrodes [18]. Rectal temperature was maintained at 37°C ± 0.5°C. For extracellular recordings, 75 μm diameter stainless steel electrodes, insulated except for their cut tips, were used. Recordings were obtained by stimulating MPP inputs to the DG molecular layer and separately the CA2/3 inputs to the SO of CA1. Two stimulating electrodes were implanted in the right hemisphere (RHS) to evoke field potentials in their respective strata. One was positioned to activate medial perforant path (MPP) inputs running in the angular bundle (0.6 mm anterior to lambda, 4 mm lateral to bregma, 2.6 mm depth from dura). The other electrode was placed in the CA1 SO (-3.3 mm posterior, 1.8 mm lateral to bregma, 2.5 mm depth from dura). Recording electrodes were placed ipsilaterally in the right hemisphere DG (3.8 mm posterior, 2.5 mm lateral to bregma, 2.8 mm depth from dura) and contralaterally in the CA1 SO (3.3 mm posterior, 1.8 mm lateral to bregma, 1.7 mm depth from dura). Final coordinates were adjusted to optimize the field potential recordingss. Post-mortem histological confirmation was performed for CA1 SO placements (**Fig. S1D**), while the stimulation of MPP inputs was confirmed electrophysiologically via paired-pulse depression (**Fig. S1C**).

Signals were amplified and filtered (0.1 Hz - 3.0 kHz). Baseline stimulation intensities were set such that MMP inputs evoked a field potential population spike response of ≥2 mV, and in SO, a fEPSP response amplitude of 0.2-0.5 mV. Baseline recordings (0.03 Hz diphasic pulses, 150 μs per half-wave, 10 min) were followed by SO TBS priming (4 trains, each containing 5 bursts at 5 Hz, each burst 5 pulses at 200 Hz, 150 μs pulses). Test pulses resumed 5-30 min before LTP induction in the DG MML by delta-burst stimulation (6 trains, each train 5 bursts at 1 Hz, each burst 10 pulses at 400 Hz, 250 μs pulses). Responses were followed for 120 min post-LTP induction.

### Whole-cell astrocyte patch-clamp recordings

Slices were incubated with 0.5 μM sulforhodamine 101 (SR101, Sigma S7635) at 32.5°C for 30 min for astrocyte labeling, followed by incubating them at room temperature in ACSF for at least 60 min before transferring them to the recording chamber. Hippocampal slices were submerged in ACSF (2.5 ml/min) and the SR101-labeled astrocytes in DG MML were identified in epifluorescence mode via green LED excitation using an Olympus URFLT microscope (Olympus BX51WI, 10x and 40x objective, **Fig. S2A**,**B**) and were patch-clamped under differential interference contrast (DIC, **Fig. S2B**) mode using borosilicate micropipettes (2.5-4.5 MΩ, Sutter Instrument Co.) filled with intracellular solution (in mM: 130 KMeSO_4_, 10 HEPES, 4 Na_2_ATP, 0.4 Na_2_GTP, 10 Na_2_phosphocreatine,4 MgCl_2_, pH 7.2, 295-300 mOsM). Calcium clamp experiments were conducted by using the intracellular solution supplemented with 0.45 mM ethylene glycol-bis (β-aminoethyl ether)-N, N, N′, N′-tetra-acetic acid (EGTA, Sigma 67-42-5) and 0.14 mM CaCl_2_ (yielding 50-80 nM free Ca^2+^). Astrocyte identity was confirmed by a low resting membrane potential (V_m_ ≥ -80 mV), and linear current-voltage responses (**Fig. S2C**). Programmable stimulators were used to stimulate fibers in CA1 SO and DG MML as described above. Synaptic potentials recorded from the patched astrocytes (AfEPSPs) were smaller than typical extracellularly recorded fEPSPs (**Fig. S2D**). To facilitate LTP induction in the MML under EGTA-Ca^2+^-clamped conditions, 100 μM glycine in combination with 0.2 μM gabazine was bath-applied throughout the experiments [19]. To qualitatively confirm MPP stimulation in the DG, paired-pulse tests were performed by delivering two pulses 50 ms apart (**Fig. S2E**).

### Calcium imaging of astrocytes

Astrocytic calcium dynamics were investigated in horizontal hippocampal slices (300 μm) from Aldh1l1-cre ERT2 ^+/-^ x Ai148 GCaMP6f (TIT2L-GC6f-ICL-tTA2)^+/-^ or Aldh1l1-cre ERT2 ^+/-^ x Ai148 GCaMP6f (TIT2L-GC6f-ICL-tTA2)^+/+^ mice. Slices, maintained at 32°C in a perfusion chamber with 2-3 mL/min ACSF flow, were imaged using a Nikon C2 confocal microscope (16x water immersion objective, NA = 0.8, with 2x optical zoom). The focal plane was consistently positioned ≥ 40 μm below the slice surface, and optical zoom was applied to target dendritic regions of interest. GCaMP6f fluorescence was excited with a 488 nm Argon laser (laser power: 5%; gain: 50), and emitted light was collected using a 505-520 nm filter.

Baseline calcium fluorescence was recorded at 1 frame/sec for three 2-minute epochs over a 10-minute period. Slices then received either 2xTBS priming stimulation in SO or not, and subsequent calcium responses were recorded from astrocytes in the molecular layer of the DG during and immediately after priming (three 2-min epochs across 10 min) and again beginning 20 min post-priming in CA1.

In a separate set of experiments, receptor inhibitors were introduced after a 10-minute baseline recording. Following a 10-minute equilibration period to allow for the inhibitors to enter the slice, a new baseline was imaged in the presence of inhibitors to assess if the drugs themselves had any effect on baseline calcium events in astrocytes. Subsequent epochs were then imaged in the presence or absence of priming. After a 20-minute wash-out period, the final 10 minutes of imaging consisted of three 2-minute epochs without inhibitors.

### Drugs

MRS 2719 (Cat. No. 2157) and pirenzepine dihydrochloride (Cat. No. 1071) were obtained from Tocris Bioscience. Kynurenic acid (K3375) and atropine (A0132) were obtained from Merck Millipore. (R, S)-MCPG (ab120033) was obtained from Abcam, and D-AP5 (HB0225) and gabazine (SR 95531 hydrobromide, HB 901) from Hello Bio. LY3020371 (cat. NO. HY-123820) was obtained from MedChemExpress. Glycine was obtained from Sigma (Sigma, G7126). Except for (R, S)-MCPG and LY3020371, all drugs were dissolved in Milli-Q water. (R, S)-MCPG was dissolved in 0.1 M NaOH, and LY3020371 was dissolved in DMSO. All drugs were diluted at least 1000-fold in ACSF before bath application.

For pharmacological priming experiments, recombinant rat TNF protein (R&D Systems, 510-RT) was reconstituted at 10 μg/mL in filter-sterilized PBS containing 0.1% bovine serum albumin (BSA) as a vehicle. LY354740 (Cat. No. 3246), a highly selective agonist of group II mGlu receptors, was obtained from Tocris and dissolved using 0.1 M NaOH. 77-LH-28-1 oxalate (560085-12-3), an agonist of M1 muscarinic acetylcholine receptors, was obtained from Aobious and was dissolved in DMSO. Across all treatment groups, for control experiments the corresponding drug vehicle was bath-applied.

### XPro1595 administration

To investigate the role of soluble TNF (sTNF) acting on TNFR1 in mediating metaplasticity, Sprague-Dawley rats or WT C57BL/6 mice were administered either saline vehicle control or XPro1595 (10 mg/kg, s.c.), 12-16 h before undertaking fEPSP recordings. XPro1595 (kindly provided by INmuneBio), is a PEGylated human TNF variant that selectively sequesters sTNF by forming heterotrimers, effectively preventing sTNF from binding to TNFR1, while leaving membrane-bound TNF signaling intact [20].

### Data analysis

For both *in vivo* and *in vitro* field potential recordings, fEPSP slopes were analyzed using custom LabVIEW software. Baseline values were calculated as the average slope for 10 min preceding LTP induction. Data were normalized to baseline and expressed as percentage change. LTP was calculated from the final 10 min of recording.

For whole-cell clamped astrocytes, data were acquired using Clampex version 10.7 and analysed using Clampfit version 10.7 (Molecular Devices). The initial slopes of the fEPSPs were measured offline and expressed as a percentage of the baseline value obtained by averaging the fEPSP slope for the 10 min preceding TBS.

Astrocyte Ca^2+^ event analysis was performed utilizing GCaMP6f fluorescence recordings quantified via the open-source AQUA (Astrocyte Quantitative Analysis) plugin in ImageJ (Fiji) [21]. Within each time-series image stack, regions of interest (ROIs) were manually delineated around individual astrocytes, and Ca^2+^ events were detected based on the AQUA plugin’s default settings, incorporating spatiotemporal filters for robust signal detection above noise. Event specificity was ensured by applying an activity threshold typically set at 2-5 σ (standard deviations above baseline), requiring a minimum duration of approximately 2 to 4 frames, and a minimum area of about 8 pixels to Ca^2+^ events, thereby excluding transient noise and single-pixel fluctuations. The resulting events were summed across 2-minute epochs for experimental conditions, including baseline, the presence (Primed, n=18 slices) and absence (Control, n=16 slices) of priming, and 20-30 min post-priming. Results were expressed as a percentage change from baseline. A similar analysis was performed for experiments conducted in the presence of a cocktail of inhibitors under control (n=7 slices) and SO priming conditions (n=9 slices).

All data are expressed as mean ± SEM. Throughout all the experiments, statistical comparisons (unpaired or paired Student’s t-test, one-way ANOVA, or two-way ANOVA with appropriate post-hoc tests) were performed using GraphPad Prism 10.6. Significance was set at *p* < 0.05.

## Acknowledgments

This work was supported by grants from the Health Research Council of New Zealand (Grant No. 18/245 to W.C.A. and Grant No. 22/177 to W.C.A. and S.S.) and Canadian Institute of Health Research (CIHR) and Natural Sciences and Engineering Research Council of Canada (NSERC) for the work undertaken at McGill University. S.S. was supported by a New Zealand International Doctoral Scholarship, a Roche Hanns Möhler Doctoral Scholarship, and a Brain Research New Zealand research grant. We thank David Stellwagen’s laboratory for providing a supportive research environment and Hooman Salahi (McGill University, Montreal) for performing the genotyping of the samples used in the mouse experiments. We sincerely thank C. J. Barnum (INmuneBio) for kindly providing XPro1595.

## References

1. Abraham, W.C., O.D. Jones, and D.L. Glanzman, Is plasticity of synapses the mechanism of long-term memory storage? NPJ Sci Learn, 2019. 4: p. 9.

2. Abraham, W.C. and M.F. Bear, Metaplasticity: the plasticity of synaptic plasticity. Trends Neurosci, 1996. 19(4): p. 126–30.

3. Abraham, W.C., Metaplasticity: tuning synapses and networks for plasticity. Nat Rev Neurosci, 2008. 9(5): p. 387.

4. Bienenstock, E.L., L.N. Cooper, and P.W. Munro, Theory for the development of neuron selectivity: orientation specificity and binocular interaction in visual cortex. J Neurosci, 1982. 2(1): p. 32–48.

5. Hulme, S.R., et al., Calcium-dependent but action potential-independent BCM-like metaplasticity in the hippocampus. J Neurosci, 2012. 32(20): p. 6785–94.

6. Singh, A., et al., Pathway-specific TNF-mediated metaplasticity in hippocampal area CA1. Scientific Reports, 2022. 12(1): p. 1746.

7. Jones, O.D., et al., Astrocyte Ca<sup>2+</sup> signalling mediates long-distance metaplasticity in the hippocampal CA1. bioRxiv, 2023: p. 2023.06.19.545623.

8. Jones, O.D., S.R. Hulme, and W.C. Abraham, Purinergic receptor- and gap junction-mediated intercellular signalling as a mechanism of heterosynaptic metaplasticity. Neurobiol Learn Mem, 2013. 105: p. 31–9.

9. Konietzko, U. and C.M. Muller, Astrocytic dye coupling in rat hippocampus: topography, developmental onset, and modulation by protein kinase C. Hippocampus, 1994. 4(3): p. 297–306.

10. Habbas, S., et al., Neuroinflammatory TNFalpha Impairs Memory via Astrocyte Signaling. Cell, 2015. 163(7): p. 1730–41.

11. Jourdain, P., et al., Glutamate exocytosis from astrocytes controls synaptic strength. Nat Neurosci, 2007. 10(3): p. 331–9.

12. Domercq, M., et al., P2Y1 receptor-evoked glutamate exocytosis from astrocytes: control by tumor necrosis factor-alpha and prostaglandins. J Biol Chem, 2006. 281(41): p. 30684–96.

13. Madisen, L., et al., A robust and high-throughput Cre reporting and characterization system for the whole mouse brain. Nature Neuroscience, 2010. 13(1): p. 133–140.

14. Ohline, S.M., et al., Egr1 Expression Is Correlated With Synaptic Activity but Not Intrinsic Membrane Properties in Mouse Adult-Born Dentate Granule Cells. Hippocampus, 2024. 34(12): p. 729–743.

15. Heir, R., et al., Astrocytes Are the Source of TNF Mediating Homeostatic Synaptic Plasticity. J Neurosci, 2024. 44(14).

16. Petersen, R.P., et al., Electrophysiological identification of medial and lateral perforant path inputs to the dentate gyrus. Neuroscience, 2013. 252: p. 154–68.

17. Abraham, W.C. and N. McNaughton, Differences in synaptic transmission between medial and lateral components of the perforant path. Brain Res, 1984. 303(2): p. 251–60.

18. Christie, B.R. and W.C. Abraham, Priming of associative long-term depression in the dentate gyrus by θ frequency synaptic activity. Neuron, 1992. 9(1): p. 79–84.

19. Sateesh, S. and W.C. Abraham, Differential Astrocyte-Supplied NMDAR Co-Agonist for CA1 Versus Dentate Gyrus Long-Term Potentiation. Hippocampus., 2025. 35(5).

20. MacPherson, K.P., et al., Peripheral administration of the soluble TNF inhibitor XPro1595 modifies brain immune cell profiles, decreases beta-amyloid plaque load, and rescues impaired long-term potentiation in 5xFAD mice. Neurobiol Dis, 2017. 102: p. 81–95.

21. Wang, Y., et al., Accurate quantification of astrocyte and neurotransmitter fluorescence dynamics for single-cell and population-level physiology. Nat Neurosci, 2019. 22(11): p. 1936–1944.

22. Sherwood, M.W., et al., Astrocytic IP(3) Rs: Contribution to Ca(2+) signalling and hippocampal LTP. Glia, 2017. 65(3): p. 502–513.

23. Sherwood, M.W., et al., Astrocytic IP(3)Rs: Beyond IP(3)R2. Front Cell Neurosci, 2021. 15: p. 695817.

24. Bukalo, O., et al., Synaptic plasticity by antidromic firing during hippocampal network oscillations. Proc Natl Acad Sci U S A, 2013. 110(13): p. 5175–80.

25. Araque, A., et al., Synaptically released acetylcholine evokes Ca2+ elevations in astrocytes in hippocampal slices. J Neurosci, 2002. 22(7): p. 2443–50.

26. Walker, A.G., et al., Co-Activation of Metabotropic Glutamate Receptor 3 and Beta-Adrenergic Receptors Modulates Cyclic-AMP and Long-Term Potentiation, and Disrupts Memory Reconsolidation. Neuropsychopharmacology, 2017. 42(13): p. 2553–2566.

27. Zur Nieden, R. and J.W. Deitmer, The Role of Metabotropic Glutamate Receptors for the Generation of Calcium Oscillations in Rat Hippocampal Astrocytes In Situ. Cerebral Cortex, 2005. 16(5): p. 676–687.

28. Wu, Y.-W., et al., Spatiotemporal calcium dynamics in single astrocytes and its modulation by neuronal activity. Cell Calcium, 2014. 55(2): p. 119–129.

29. Cotrina, M.L., et al., ATP-mediated glia signaling. J Neurosci, 2000. 20(8): p. 2835–44.

30. Shigetomi, E., et al., Role of Purinergic Receptor P2Y1 in Spatiotemporal Ca(2+) Dynamics in Astrocytes. J Neurosci, 2018. 38(6): p. 1383–1395.

31. Bowser, D.N. and B.S. Khakh, ATP excites interneurons and astrocytes to increase synaptic inhibition in neuronal networks. J Neurosci, 2004. 24(39): p. 8606–20.

32. Orellana, J.A., et al., ATP and glutamate released via astroglial connexin 43 hemichannels mediate neuronal death through activation of pannexin 1 hemichannels. J Neurochem, 2011. 118(5): p. 826–40.

33. Singh, A., et al., Tumor Necrosis Factor-alpha-Mediated Metaplastic Inhibition of LTP Is Constitutively Engaged in an Alzheimer’s Disease Model. J Neurosci, 2019. 39(46): p. 9083–9097.

34. Park, K.M. and W.J. Bowers, Tumor necrosis factor-alpha mediated signaling in neuronal homeostasis and dysfunction. Cell Signal, 2010. 22(7): p. 977–83.

35. Nikolic, L., et al., Blocking TNFalpha-driven astrocyte purinergic signaling restores normal synaptic activity during epileptogenesis. Glia, 2018. 66(12): p. 2673–2683.

36. Liu, D.D., Q. Yang, and S.T. Li, Activation of extrasynaptic NMDA receptors induces LTD in rat hippocampal CA1 neurons. Brain Res Bull, 2013. 93: p. 10–6.

37. Bushong, E.A., et al., Protoplasmic astrocytes in CA1 stratum radiatum occupy separate anatomical domains. J Neurosci, 2002. 22(1): p. 183–92.

38. Perea, G., M. Navarrete, and A. Araque, Tripartite synapses: astrocytes process and control synaptic information. Trends Neurosci, 2009. 32(8): p. 421–31.

39. Hulme, S.R., et al., Mechanisms of heterosynaptic metaplasticity. Philos Trans R Soc Lond B Biol Sci, 2014. 369(1633): p. 20130148.

40. Pinto-Duarte, A., et al., Impairments in remote memory caused by the lack of Type 2 IP(3) receptors. Glia, 2019. 67(10): p. 1976–1989.

41. Perea, G. and A. Araque, Properties of synaptically evoked astrocyte calcium signal reveal synaptic information processing by astrocytes. J Neurosci, 2005. 25(9): p. 2192–203.

42. Miguel-Quesada, C., et al., Astrocytes adjust the dynamic range of cortical network activity to control modality-specific sensory information processing. Cell Reports, 2023. 42(8): p. 112950.

43. Zeng, W., et al., A new mode of Ca2+ signaling by G protein-coupled receptors: gating of IP3 receptor Ca2+ release channels by Gbetagamma. Curr Biol, 2003. 13(10): p. 872–6.

44. Haustein, Martin D., et al., Conditions and Constraints for Astrocyte Calcium Signaling in the Hippocampal Mossy Fiber Pathway. Neuron, 2014. 82(2): p. 413–429.

45. Latour, I., et al., Differential mechanisms of Ca2+ responses in glial cells evoked by exogenous and endogenous glutamate in rat hippocampus. Hippocampus, 2001. 11(2): p. 132–45.

46. Murphy, S., B. Pearce, and C. Morrow, Astrocytes have both M1 and M2 muscarinic receptor subtypes. Brain Res, 1986. 364(1): p. 177–80.

47. Shelton, M.K. and K.D. McCarthy, Hippocampal astrocytes exhibit Ca2+-elevating muscarinic cholinergic and histaminergic receptors in situ. J Neurochem, 2000. 74(2): p. 555–63.

48. Navarrete, M., et al., Astrocytes mediate in vivo cholinergic-induced synaptic plasticity. PLoS Biol, 2012. 10(2): p. e1001259.

49. Takata, N., et al., Astrocyte calcium signaling transforms cholinergic modulation to cortical plasticity in vivo. J Neurosci, 2011. 31(49): p. 18155–65.

50. Lee, M.-C., R. Yasuda, and M.D. Ehlers, Metaplasticity at Single Glutamatergic Synapses. Neuron, 2010. 66(6): p. 859–870.

51. Yang, Q., et al., Hippocampal synaptic metaplasticity requires the activation of NR2B-containing NMDA receptors. Brain Res Bull, 2011. 84(2): p. 137–43.

52. Amaral, D.G. and M.P. Witter, The three-dimensional organization of the hippocampal formation: a review of anatomical data. Neuroscience, 1989. 31(3): p. 571–91.

53. Yassa, M.A. and C.E. Stark, Pattern separation in the hippocampus. Trends Neurosci, 2011. 34(10): p. 515–25.

54. Neunuebel, Joshua P. and James J. Knierim, CA3 Retrieves Coherent Representations from Degraded Input: Direct Evidence for CA3 Pattern Completion and Dentate Gyrus Pattern Separation. Neuron, 2014. 81(2): p. 416–427.

55. Atucha, E., et al., Recalling gist memory depends on CA1 hippocampal neurons for lifetime retention and CA3 neurons for memory precision. Cell Reports, 2023. 42(11): p. 113317.

56. Miller, E.K. and J.D. Cohen, An integrative theory of prefrontal cortex function. Annu Rev Neurosci, 2001. 24: p. 167–202.

57. D’Esposito, M. and B.R. Postle, The cognitive neuroscience of working memory. Annu Rev Psychol, 2015. 66: p. 115–42.

58. Abraham, W.C. and A. Robins, Memory retention--the synaptic stability versus plasticity dilemma. Trends Neurosci, 2005. 28(2): p. 73–8.

59. Zenke, F. and W. Gerstner, Hebbian plasticity requires compensatory processes on multiple timescales. Philos Trans R Soc Lond B Biol Sci, 2017. 372(1715).

60. Thompson, A., et al., Brain-wide circuit-specific targeting of astrocytes. Cell Rep Methods, 2023. 3(12): p. 100653.

61. Wang, X., et al., Astrocytic Ca2+ signaling evoked by sensory stimulation in vivo. Nat Neurosci, 2006. 9(6): p. 816–23.

62. Fillit, H., et al., Elevated circulating tumor necrosis factor levels in Alzheimer’s disease. Neurosci Lett, 1991. 129(2): p. 318–20.

63. McCoy, M.K. and M.G. Tansey, TNF signaling inhibition in the CNS: implications for normal brain function and neurodegenerative disease. J Neuroinflammation, 2008. 5: p. 45.

64. Delcy, S.A.S., et al., Microglia depletion improves hippocampal circuit function after mild traumatic brain injury in male mice. Brain Behav Immun, 2026. 131: p. 106178.

65. Macek, T.A., et al., Differential involvement of group II and group III mGluRs as autoreceptors at lateral and medial perforant path synapses. J Neurophysiol, 1996. 76(6): p. 3798–806

