## Supplementary figures for "Transregional astrocyte-dependent metaplasticity in the hippocampus"

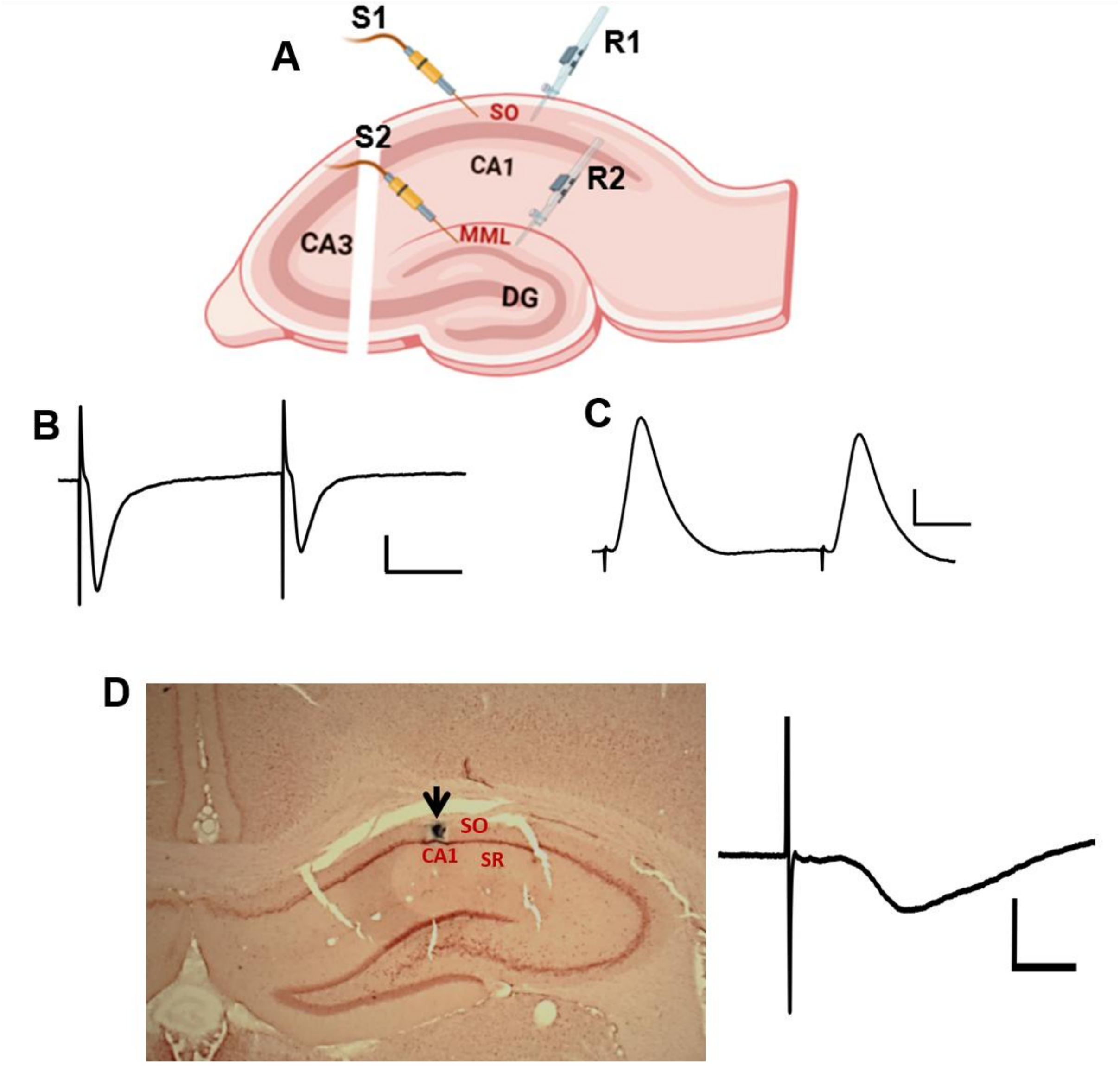


**Figure S1: (A)** Schematic electrode placements in stratum oriens (SO) and middle molecular layer (MML) of the dentate gyrus (DG) in an acute transverse rat hippocampal slice. Priming stimulation was delivered via a stimulating electrode (S1) in SO. LTP was delivered 30 min later via a second stimulating electrode placed in the MML (S2). Recording electrode: R1 recorded the change in synaptic efficacies in the primed pathway (SO), and R2 recorded the changes to LTP-inducing stimulus in the test pathway (MML). **(B,C)** Representative waveform average of 5 sweeps showing paired-pulse depression (PPD) recorded in the MML *in vitro* **(B)** and in the dentate hilus *in vivo* **(C)**. Two pulses were delivered at a 50-ms inter-pulse interval to the medial perforant path (MPP)-DG fibres running in the MML. Scale bars: 0.5 mV, 20 ms **(B)**, 2 mV, 10 ms **(C)**. **(D)** Representative hippocampal section confirming the *in vivo* electrode placement in SO using lesion stain (black circle indicated by arrow) with Prussian blue and neutral red. ***Inset:*** Representative waveform of fEPSP evoked by stimulating afferents and recording from a group of stratum oreins (SO) synapses *in vivo*. Scale bar: 0.5 mV, 5 ms.

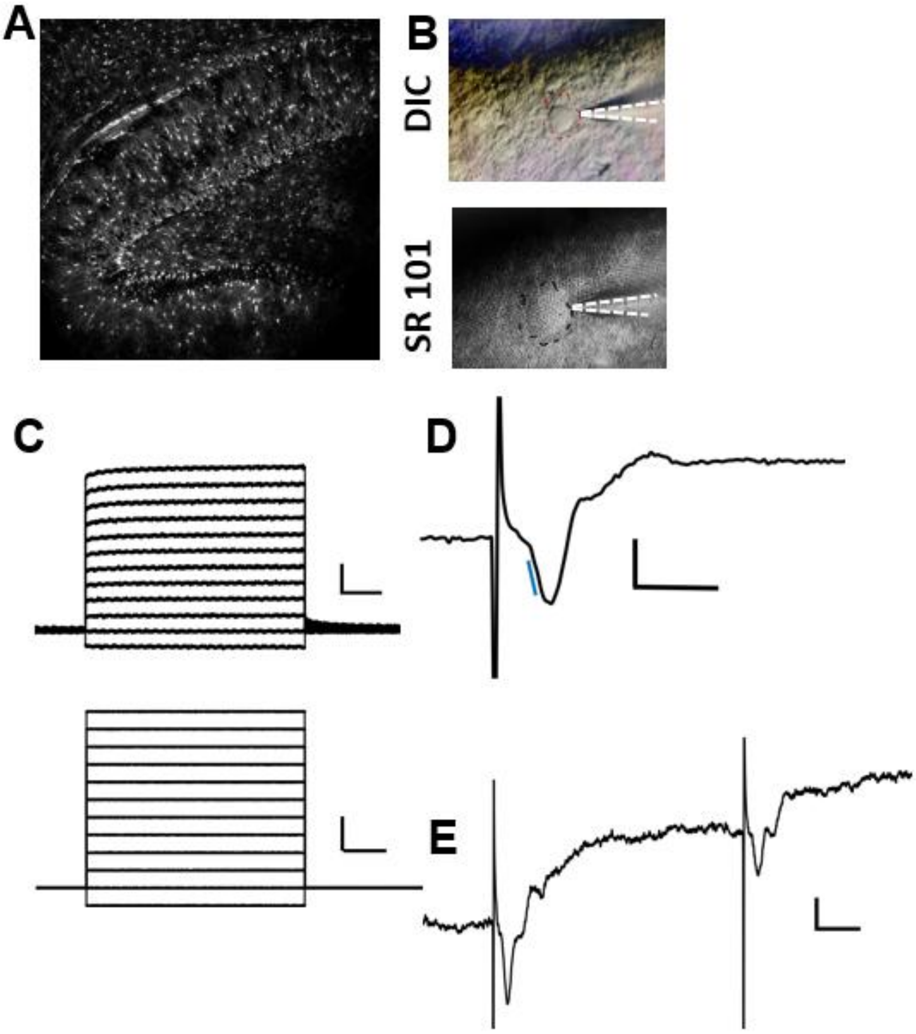


**Figure S2: (A)** Representative confocal image of astrocytes labelled with SR101 using a 10x water immersion objective. **(B)** Representative images showing differential interference contrast (DIC) and the corresponding fluorescence image of an SR 101 patched astrocyte in MML, with a glass pipette shown in a white dotted line, using a 40x water-immersion objective lens. (**C**) Bottom, schematic of 200 pA current steps used to generate $V_m$ deflections, and Top, the corresponding traces of current-voltage relation from a single whole-cell patch-clamped astrocyte. Scale bars: 2 mV, 100 ms; 200 pA, 100 ms. **(D)** A representative evoked field excitatory postsynaptic potential (fEPSP) was recorded via a patched astrocyte in MML (AfEPSP) following test-pulse stimulation of the MPP inputs. Blue line indicates the slope measurement. Scale bar: 1 mV, 5 ms. **(E)** Representative average waveform (4 sweeps) of evoked synaptic field potentials recorded through the astrocyte membrane in response to paired-pulse stimulation at a 50-ms inter-pulse interval, showing paired-pulse depression at MPP-DG synapses. Scale bar: 0.5 mV, 10 ms.

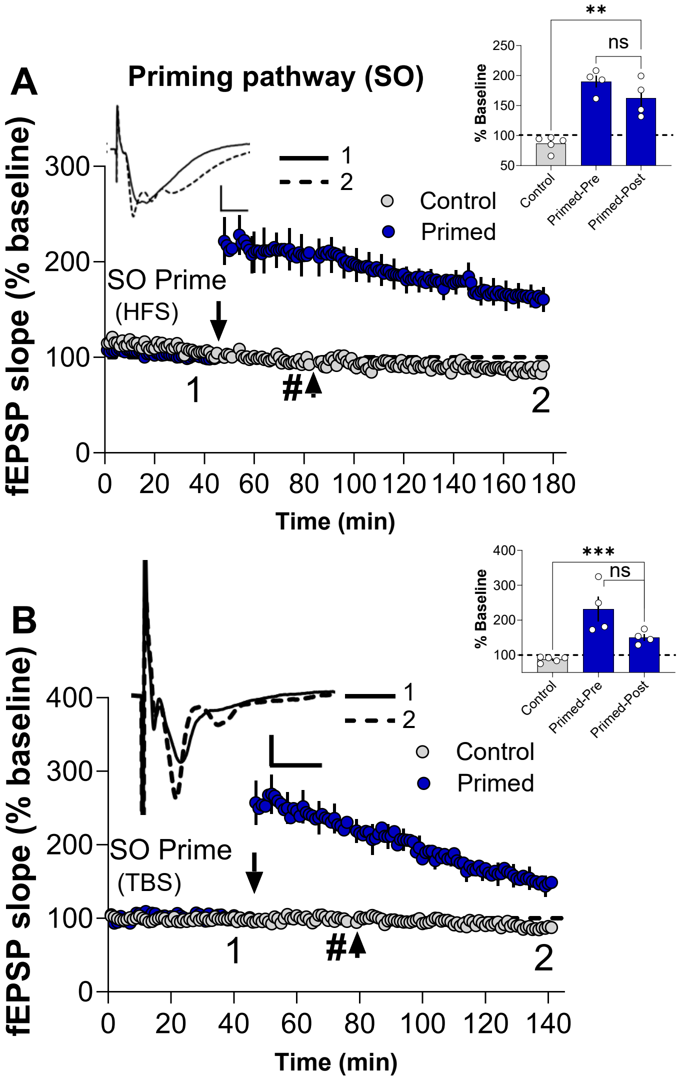


**Figure S3: Recordings from priming pathway stratum oriens (SO) with and without priming stimulation for experiments shown in (Fig. 1A,B).**

**(A)** HFS priming stimulation of afferents in SO resulted in homosynaptic LTP compared to non-primed controls measured at the end of the recording period (Control: 86.8 ± 5.6%, *n* = 5; Primed-Post: 162.6 ± 15.4%, *n* = 4, $t_{(7)}$ = 5.1, *p* = 0.0015). The magnitude of SO LTP measured immediately before the conditioning stimulus was delivered to MML (Primed-Pre (#):190.1 ± 10.1%, *n* = 4) was found not to be significantly different from the LTP measured at the end of the experiment (*paired* $t_{(3)}$ = 2.1, *p* = 0.12). **(B)**TBS priming also resulted in substantial LTP compared to non-primed controls, measured at the end of 90 min recording period (Control: 87.5 ± 3.6%, *n* = 5; Primed-Post: 150.5 ± 9.5%, *n* = 4, $t_{(7)}$ = 6.8, *p* =0.0003). Similar to previous observation, SO LTP measured immediately before conditioning stimulus in MML (Primed-Pre: 231.9 ± 35.3%) was found not different from LTP measured at the end of 90 min (*paired* $t_{(3)}$ = 2.8, *p* = 0.066). ***Inset:*** Representative waveforms are an average of 10 synaptic responses prior to SO LTP induction (**1**), and at the conclusion of the experiment (**2**). Bar graphs summarize the LTP across individual slices. Scale bars: 0.5 mV, 5 ms (**A-B**). Arrows indicate the timing of SO priming and MML LTP induction, and (**#**) indicates measurement of SO LTP 30 min post-priming or immediately before delivery of MML TBS (Primed-Pre). All data presented as mean ± SEM; ns; *p* > 0.05; ∗∗, *p* < 0.01; ∗∗∗, *p* < 0.001.

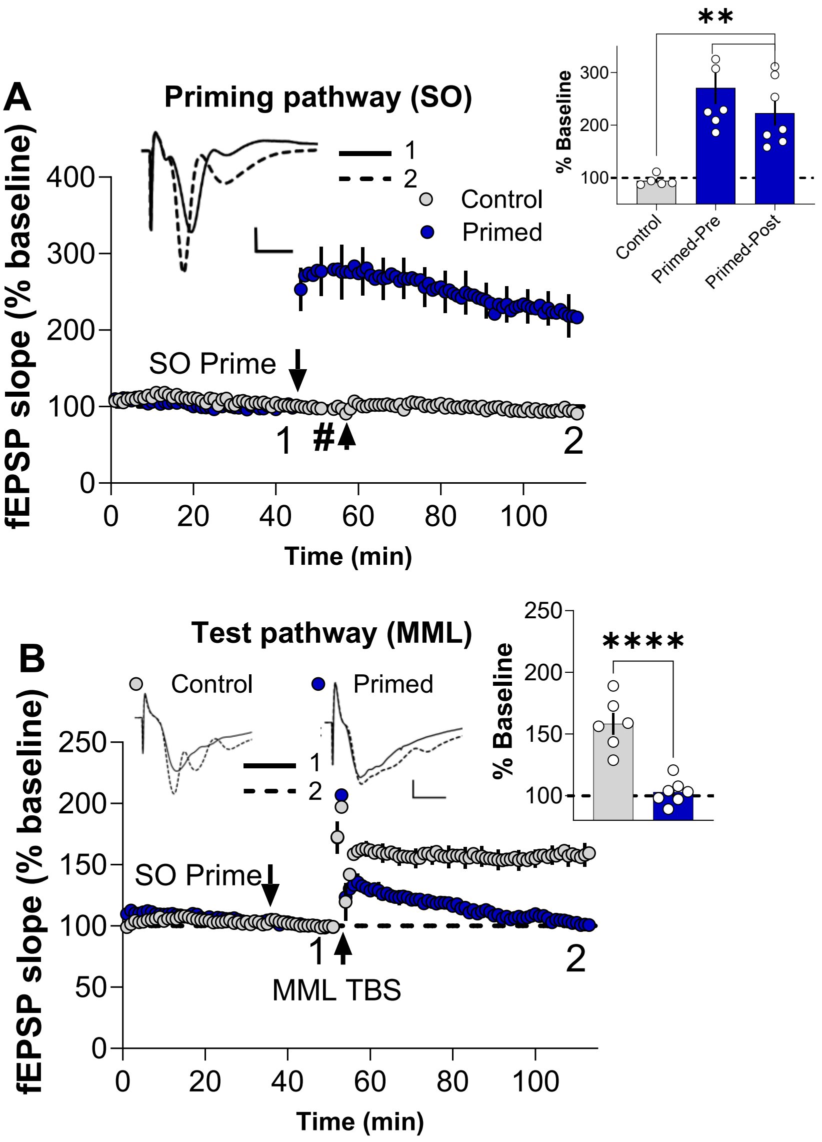


**Figure S4: TBS Priming in SO inhibits LTP in MML 5 min post-priming.**

**(A)** In the SO priming pathway, homosynaptic LTP was observed (Control: 94.3 ± 4.4%, *n* = 5; Primed-Post: 223 ± 23.8%, *n* = 7, $t_{(10)}$ = 4.5, *p* = 0.0012). The magnitude of SO LTP measured immediately before the conditioning stimulus was delivered to MML (Primed-Pre (#): 270.8 ± 30.8%, *n* = 7) was found to partially decay over the course of experimental recording (*paired* $t_{(6)}$ = 4.0, *p* = 0.0074). **(B)** In the test pathway, SO priming robustly inhibited subsequent MML LTP even with the reduced interval (Control: 158.2 ± 8.6%, *n* = 6; Primed: 102.9 ± 3.7%, *n* = 7, $t_{(11)}$ = 6.2, *p* < 0.0001). ***Inset:*** Representative waveforms are an average of 10 synaptic responses prior to SO LTP induction (**1**) and at the conclusion of the experiment (**2**). Bar graphs summarize LTP across individual experiments. Scale bars: 0.5 mV, 5 ms (**A-B**). Arrows indicate the timing of SO priming and MML LTP induction, and (**#**) indicates measurement of SO LTP 30 min post-priming or immediately before delivery of MML TBS (Primed-Pre). All data presented as mean ± SEM; ∗∗, *p* < 0.01; ∗∗∗, *p* < 0.001.

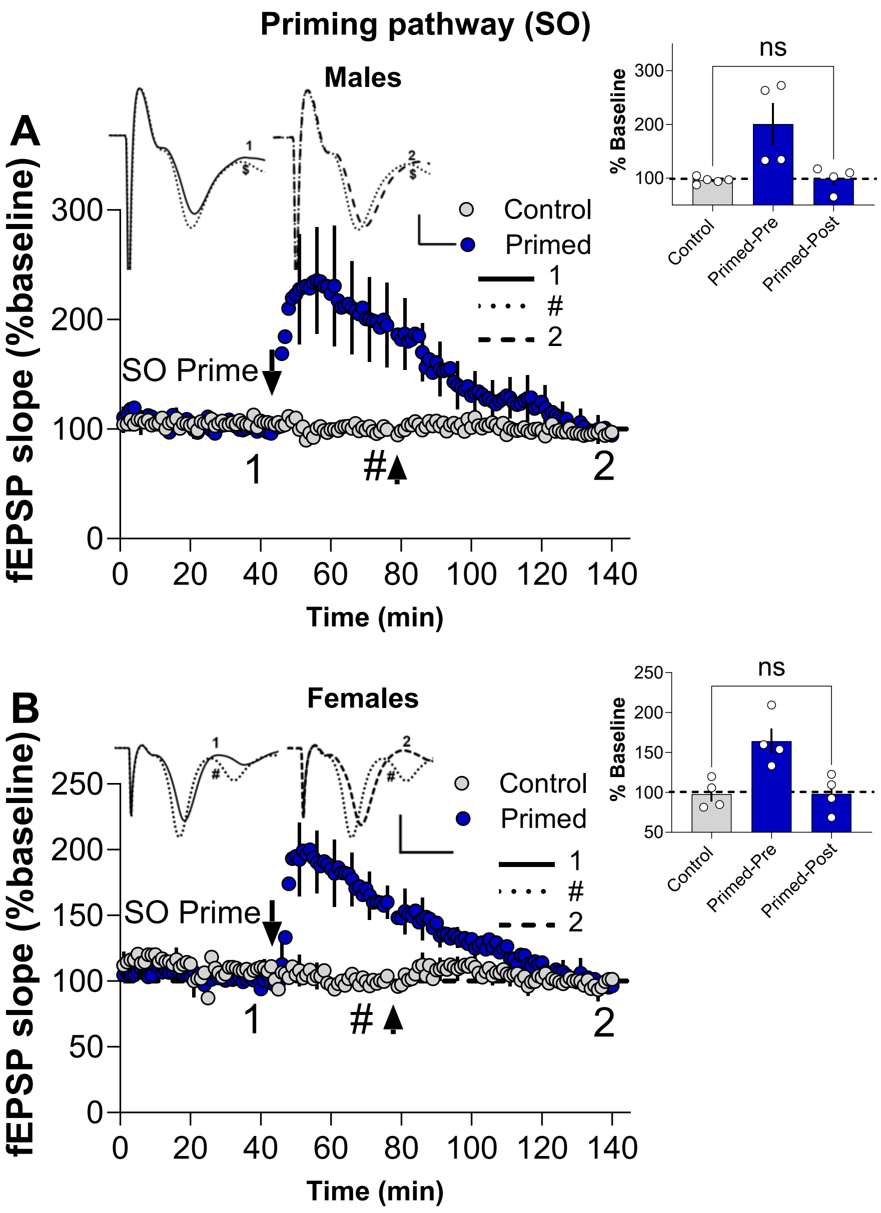


**Figure S5: Homosynaptic LTP in the SO priming pathway exhibits complete depotentiation in mouse hippocampal slices from experiments described in (Fig.1 E,F)**

Electrical priming in SO resulted in homosynaptic LTP in males **(A)** and females **(B)** that underwent complete depotentiation**.** TBS priming activity resulted in substantial potentiation as measured at 30 min post-priming in both male (Primed-Pre (#): 200.9 ± 38.8%, *n* = 4, **A**) and female hippocampus (Primed-Pre (#): 164.1 ± 16.1%, *n* = 4, **B**). In contrast to rat hippocampal recording, SO LTP in mouse hippocampus underwent complete depotentiation as measured at the end of 90 min due to the conditioning stimulus in MML, such that it was not significantly different from non-primed controls, in both **(A)** males (Control: 96.1 ± 2.7, *n* = 5; Primed-Post: 98.86 ± 11.6%, *n* = 4, $t_{(7)}$ = 0.26, *p* = 0.80) and **(B)** females (Control: 97.9.1 ± 9.1, *n* = 4; Primed-Post: 98.3 ± 11.7%, *n* = 4, $t_{(6)}$ = 2.3, *p* = 0.03, *p* = 0.98). ***Inset:*** Representative waveforms are an average of 10 synaptic responses prior to SO LTP induction (**1**), 30 min post priming (**#**), and at the conclusion of the experiment (**2**). Bar graphs summarize the LTP across individual slices. Scale bars: 0.5 mV, 5 ms (**A**); 1 mV, 5 ms (**B**). Arrows indicate the timing of SO priming and MML LTP induction, and (#) indicates measurement of SO LTP 30 min post-priming or immediately before delivery of MML TBS (Primed-Pre). All data presented as mean ± SEM; ns, *p* > 0.05.

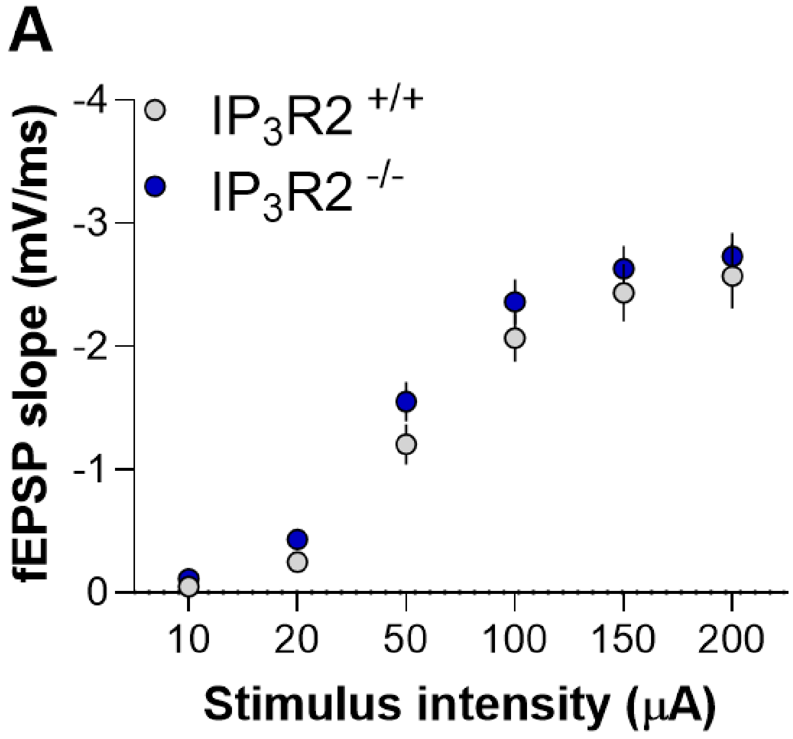


**Figure S6: fEPSP I/O curves comparing IP3R2$^{+/+}$ and IP3R2$^{-/-}$ mice.**

**(A)** Input-output (I/O) curves were generated at MML synapses of $IP_3R2^{-/-}$ and $IP_3R2^{+/+}$ littermates by delivering a series of stimulations to the MMP-DG afferent fibres using current increments (10 µA, 20 µA, 50 µA, 100 µA, 150 µA, and 200 µA) and recording the corresponding fEPSP responses. Two-way ANOVA (genotype x current intensities) indicated no significant difference in the basal synaptic transmission in $IP_3R2^{-/-}$ compared to the $IP_3R2^{+/+}$ littermates ($F_{(1, 28)}$ = 1.06, $p$ = 0.31, $IP_3R^{+/+}$: $n$ = 14; $IP_3R2^{-/-}$: $n$ = 17). All data presented as mean ± SEM.

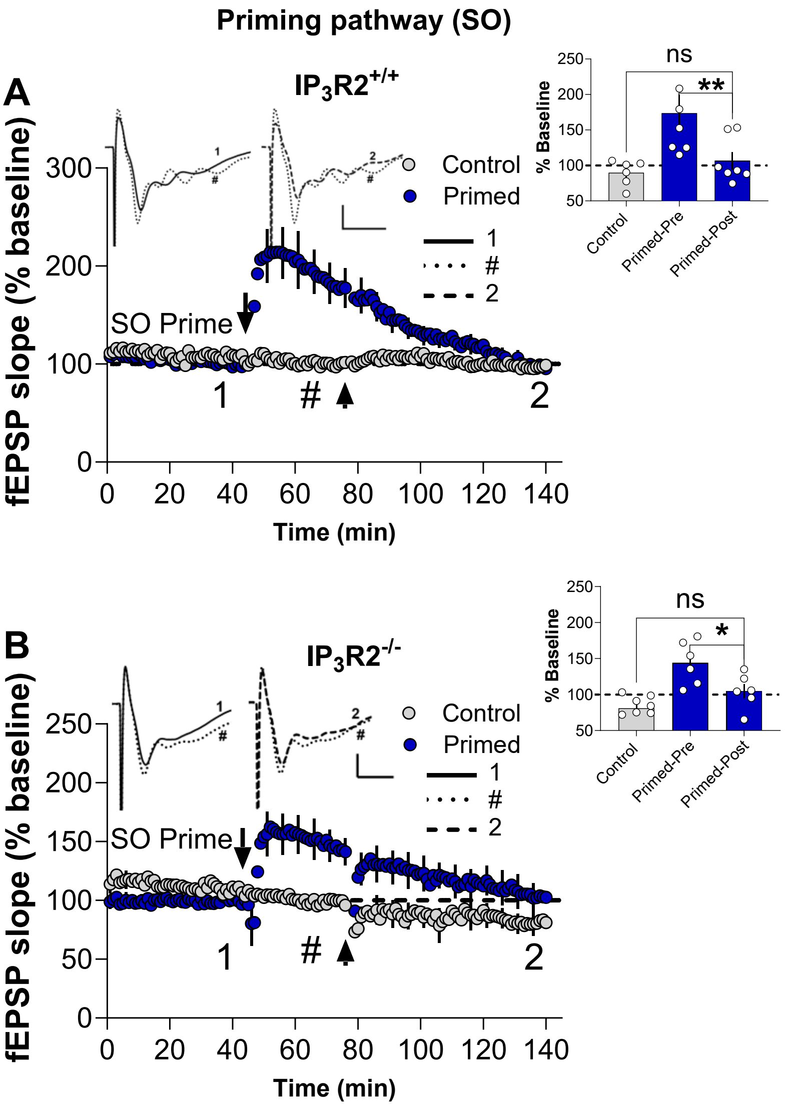


**Figure S7: SO LTP maintenance in $IP_3R2^{+/+}$ and $IP_3R2^{-/-}$mice from experiments described in (Fig.2 D,E)**

**(A,B)** TBS priming-induced homosynaptic LTP in the SO pathway undergoes depotentiation regardless of astrocytic expression of $IP_3R2$. SO priming activity resulted in substantial potentiation as measured at 30 min post-priming, both in **(A)** $IP_3R2^{+/+}$ (Primed-Pre (#): 173.6 ± 25.9%, *n* =7) and **(B)** $IP_3R2^{-/-}$ mice (Primed-Pre (#): 144.0 ± 12.3%, *n* = 6). Similar to the effect observed in the WT mouse hippocampal slices described in (**Fig. S5)**, LTP underwent complete depotentiation as measured at end of 90 min that was not significantly different to non-primed controls, in **(A)** $IP_3R2^{+/+}$ mice (Control: 89.7 ± 7.3%, *n* = 6; Primed-Post: 106.8 ± 12.1%, *n* =7, $t_{(11)}$ = 1.2, *p* = 0.27, ) and **(B)** $IP_3R2^{-/-}$ mice (Control: *n*=8, 80.9 ± 6.2%; KO Primed-Post: 104.6 ± 9.8%, *n* = 6, $t_{(12)}$ = 2.2, *p*= 0.052). Further, SO LTP compared 30 min post (Primed-Pre) to 90 min (Primed-Post) also showed a significant reduction in **(A)** $IP_3R2^{+/+}$ mice (*paired* $t_{(6)}$ = 3.9, *p* = 0.0081) and in **(B)** $IP_3R2^{-/-}$ mice (*paired* $t_{(5)}$ = 3.3, *p* = 0.0225). ***Inset:*** Representative waveforms are an average of 10 synaptic responses prior to SO LTP induction (**1**), 30 min post priming (**#**), and at the conclusion of the experiment (**2**). Bar graphs summarize the LTP across individual slices. Scale bars: 0.5 mV, 5 ms (**A-B**). Arrows indicate the timing of SO priming and MML LTP induction, and (**#**) indicates measurement of SO LTP 30 min post-priming or immediately before delivery of MML TBS (Primed-Pre). All data presented as mean ± SEM; ns, $p > 0.05$; ∗, $p < 0.05$; ∗∗, $p < 0.01$.

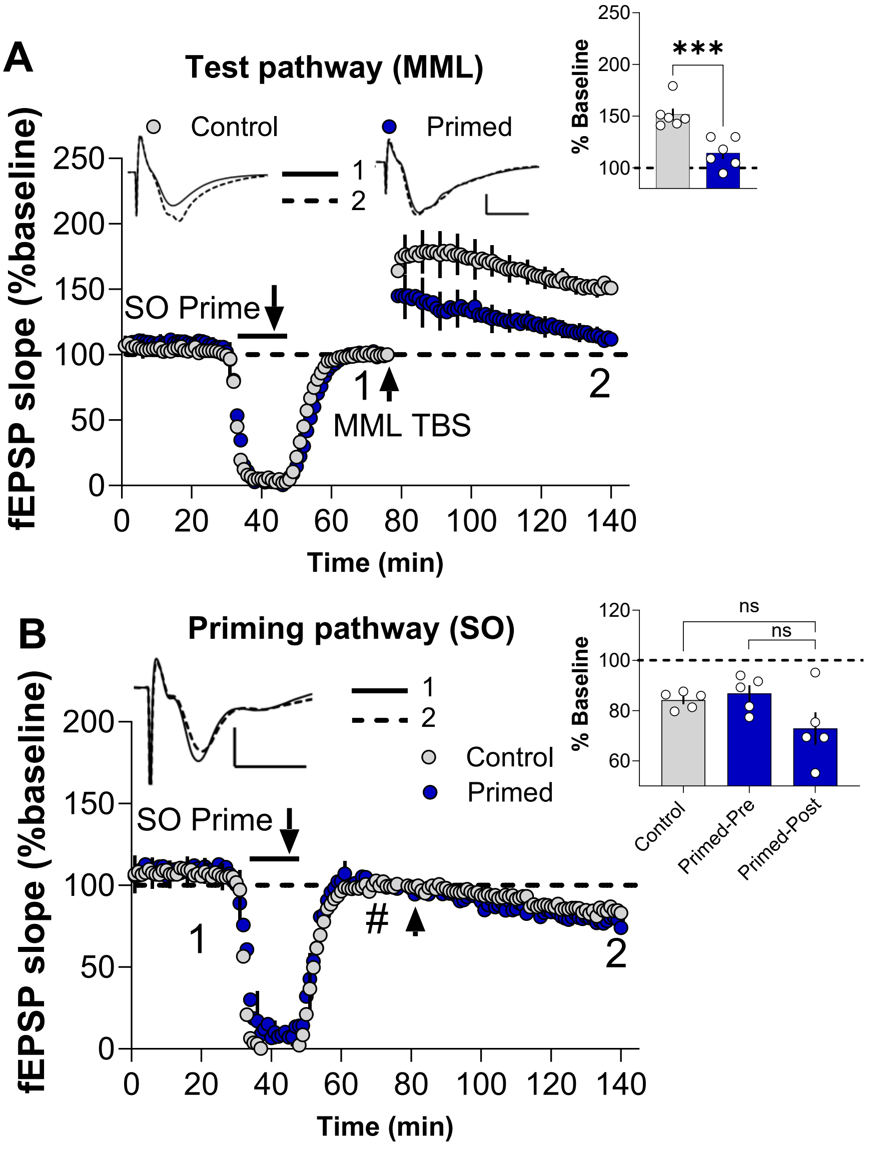


**Figure S8: Glutamatergic signalling and transregional metaplasticity**

**(A)**SO priming in the presence of a cocktail of glutamatergic inhibitors (AMPAR/KAR antagonist, 3 mM kyenurate, NMDAR antagonist, 50 μM D-AP5, and Group I/II mGluR antagonist 250 μM MCPG) did not block the transregional metaplasticity effect compared to non-primed controls (Control:151.7 ± 5.7%; *n* = 6; Primed: 114.3 ± 5.8%, *n* = 6, $t_{(10)}$ = 4.6, *p* = 0.0010). **(B)** In the priming pathway, as would be expected under conditions where NMDARs are blocked, priming stimulation did not produce any SO LTP as measured at 30 min post-priming (Primed-Pre (#): 86.8 ± 3.1%, *n* = 5, *paired* $t_{(4)}$ = 2.6, *p* = 0.058) or 90 min post (Control: 84.2 ± 1.5%, *n* = 5; Primed-Post: 72.9 ± 6.5%, $t_{(8)}$ = 1.7, *p* = 0.1290). ***Inset:*** Representative waveforms are an average of 10 synaptic responses prior to SO LTP induction (**1**) and at the conclusion of the experiment (**2**). Bar graphs summarize the LTP across individual slices. Scale bars: 1 mV, 5 ms (**A-B**). Arrows indicate the timing of SO priming and MML LTP induction, and (**#**) indicates measurement of SO LTP 30 min post-priming or immediately before delivery of MML TBS (Primed-Pre). All data presented as mean ± SEM; *ns*, *p* > 0.05; ∗∗∗*p* < 0.001.

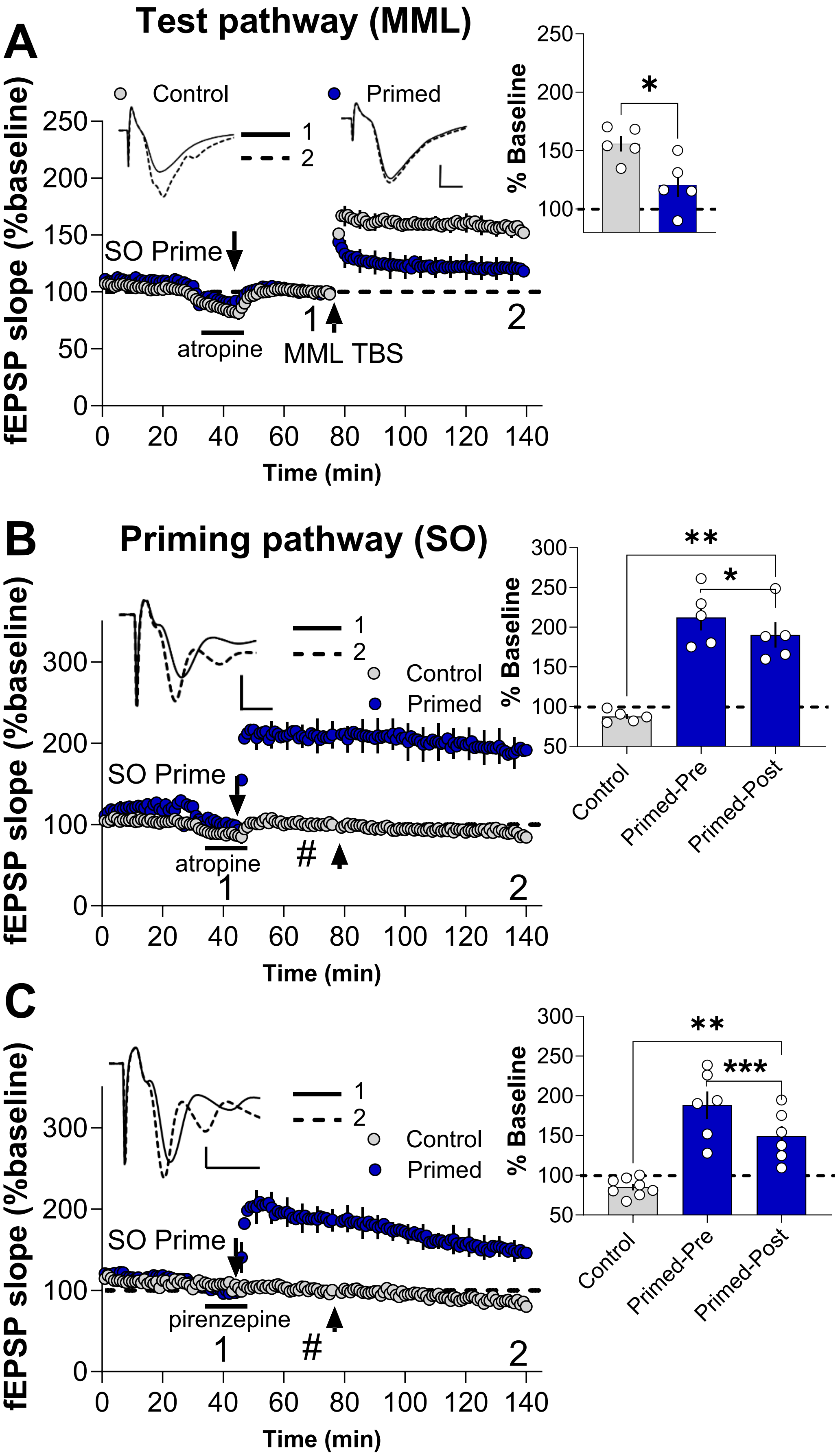
A
Test pathway (MML)
Control
Primed
1
2
SO Prime
atropine
1
MML TBS
2
fEPSP slope (%baseline)
Time (min)
% Baseline
*
B
Priming pathway (SO)
1
2
Control
Primed
SO Prime
atropine
#
1
2
fEPSP slope (%baseline)
Time (min)
% Baseline
**
*
Control
Primed-Pre
Primed-Post
C
1
2
Control
Primed
SO Prime
pirenzepine
1
#
2
fEPSP slope (%baseline)
Time (min)
% Baseline
**
***
Control
Primed-Pre
Primed-Post

**Figure S9: Cholinergic signalling and transregional metaplasticity**

(**A**) SO priming in the presence of a nonselective mAChR blocker, 10 μM atropine, significantly inhibited MML LTP compared to controls (Control: 155.9 ± 6.4%, $n$ = 5; Primed: 120.6 ± 9.9%, $n$ =5, $t_{(8)}$ =3.0, $p$ = 0.0176). **(B)** In the SO priming pathway, atropine did not affect the induction of homosynaptic LTP as measured at the end of 90 min (Control: 87.6 ± 3.2%, $n$ = 5; Primed-Post: 190.3 ± 15.7%, $n$ =5, $t_{(8)}$ = 6.4, $p$ = 0.0002) but decayed over the course of the experiment when compared 30 min and 90 min post priming (Primed-Pre: 212.1 ± 16.0%, $n$ = 5, *paired* $t_{(4)}$ = 3.9, $p$ = 0.0181). **(C)** Similar to atropine, SO homosynaptic LTP remained unaffected when TBS priming was undertaken in the presence of 20 μM pirenzepine, M1-mAChR selective antagonist (Control: 85.21 ± 4.0%, $n$ = 8; Primed-Post: 149.3 ± 13.0%, $n$ = 6, $t_{(12)}$ = 5.3, $p$ = 0.0002) that decayed over the course of the experiment when compared between 30 min and 90 min post priming (Primed-Pre (#): 188.2 ± 17.3%; $n$ = 6, *paired* $t_{(5)}$ = 7.1, $p$ = 0.0009). ***Inset:*** Representative waveforms are an average of 10 synaptic responses prior to SO LTP induction (**1**), and at the conclusion of the experiment (**2**). Bar graphs summarize the LTP across individual slices. Scale bars: 1 mV, 2 ms (**A-B**) 0.5 mV, 4 ms (**C**). Arrows indicate the timing of SO priming and MML LTP induction, and (**#**) indicates measurement of SO LTP 30 min post-priming or immediately before delivery of MML TBS (Primed-Pre). All data presented as mean ± SEM; *ns*, $p > 0.05$; ∗, $p < 0.05$; ∗∗, $p < 0.01$; ∗∗∗, $p < 0.001$.

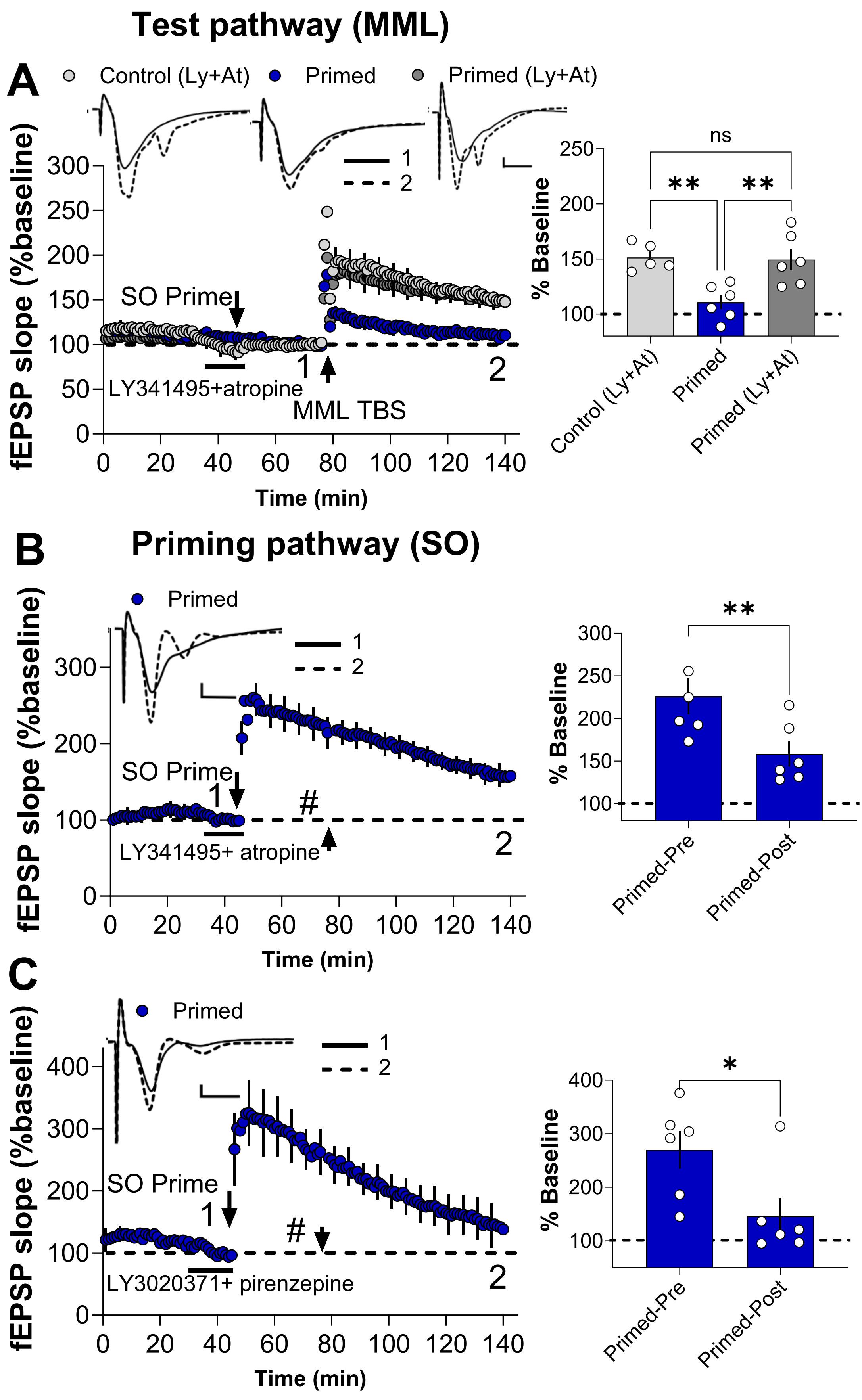

Test pathway (MML)
A
Control (Ly+At)
Primed
Primed (Ly+At)
1
2
fEPSP slope (%baseline)
SO Prime
LY341495+atropine
MML TBS
Time (min)
ns
**
**
% Baseline
Control (Ly+At)
Primed
Primed (Ly+At)
B
Priming pathway (SO)
Primed
SO Prime
#
LY341495+ atropine
Primed-Pre
Primed-Post
C
LY3020371+ pirenzepine
*

**Figure S10: Transregional metaplasticity was blocked by mAChR and mGluR antagonists**

**(A)** Priming-induced inhibition of MML LTP was blocked by non-selective mAChR and group I/II mGluR antagonists**.** One-way ANOVA indicated a significant priming effect ($F_{(2,14)}$ = 9.3, $p$ = 0.0027). SO priming delivered in the presence of a cocktail containing 100 µM LY341495, group I/II mGluR antagonist, and 10 µM atropine, a non-selective mAChR antagonist, effectively blocked the SO priming-mediated inhibition of MML LTP. The results showed that LTP levels were not significantly different from the non-primed controls (Cocktail Control no-primed: 151.4 ± 5.5%, $n$ = 5; Cocktail Primed: 149.5 ± 9.5%, $n$ = 6, post-hoc Tukey's test, $p$ = 0.98). However, SO priming in the absence of the drug significantly inhibited MML LTP (Vehicle Primed: 110.8 ± 6.4%, $n$ = *6*) when compared to LTP measured in the presence of the cocktail ($p$ = 0.0062) and in the absence of priming ($p$ = 0.0061). **(B)** SO priming resulted in substantial homosynaptic potentiation that was unaltered by the presence of 100 µM LY341495 and 10 µM atropine as measured 30 min post-priming (Primed-Pre (#): 226.0 ± 20.9%, $n$ = 6) but which decayed over the ensuing 90 min (Primed-Post: 158.6 ± 14.5%, $n$ = 6*, paired* $t_{(5)}$= 4.8, $p$ = 0.0050). **(C)** Consistent with our previous observations, SO priming-induced LTP remained unaltered by the presence of a cocktail of highly selective and potent blockers of group II mGluRs, LY3020371 (20 µM), and M1-mAChRs, pirenzepine (20 µM), as measured 30 min post-priming (Primed-Pre (#): 269.9 ± 35.4%, $n$ = 6) but which decayed over the course of the ensuing 90 min (Primed-Post: 146.0± 34.18%, $n$ = 6*, paired* $t_{(5)}$= 3.005, $p$ = 0.0299). ***Inset:*** Representative waveforms are an average of 10 synaptic responses prior to SO LTP induction (**1**) and at the conclusion of the experiment (**2**). Bar graphs summarize the LTP across individual slices. Scale bars: 0.5 mV, 5 ms (**A-C**). Arrows indicate the timing of SO priming and MML LTP induction, and (**#**) indicates measurement of SO LTP 30 min post-priming or immediately before delivery of MML TBS (Primed-Pre). All data presented as mean ± SEM; *ns*, $p$ > 0.05; $*$, $p$ < 0.05; $**$, $p$ < 0.01.

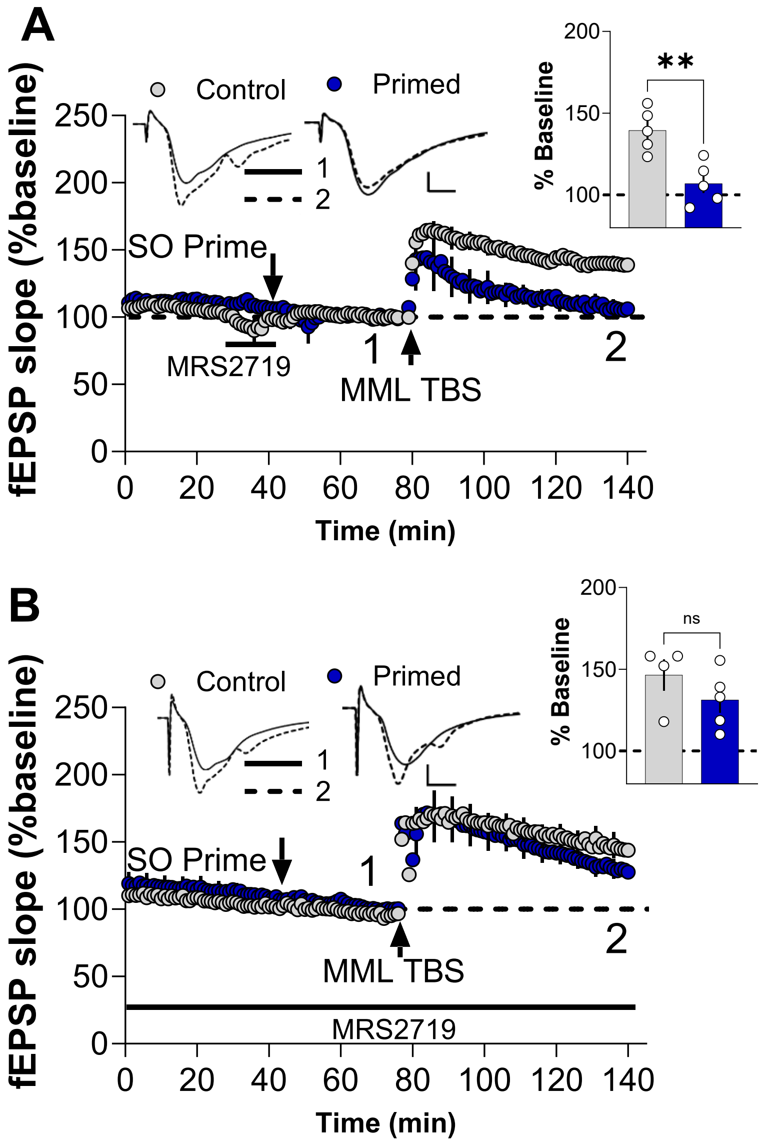


**Figure S11: Effect of the P2Y1R antagonist MRS2719 on transregional metapalsticty**

**(A)** SO priming in the presence of 10 µM MRS 2719 still significantly inhibited MML LTP (Control non-primed: 139.6 ± 5.9%, *n* =5; Primed: 107.0 ± 5.7%, *n = 5,* $t_{(8)}$ = 4.0, *p* = 0.0041). (**B**) Bath application of 10 µM MRS 2719 throughout the experimental recording restricted the priming-mediated inhibition of LTP (Control non-primed: 146.6 ± 9.6%, *n* = 4; Primed: 131.3 ± 7.9%, *n* = 5, $t_{(7)}$ = 1.2, *p* = 0.26). Note that a small run-down of baseline responses was observed in both control and primed conditions. ***Inset:*** Representative waveforms are an average of 10 synaptic responses prior to LTP induction (**1**) and at the conclusion of the experiment (**2**). Bar graphs summarize the SO priming effects in the presence of MRS 2719 on MML LTP. Scale bars: 1 mV, 2 ms (**A-B**). Arrows indicate the timing of SO priming and MML LTP induction. All data presented as mean ± SEM; *ns*, *p* > 0.05; **, *p* < 0.01.

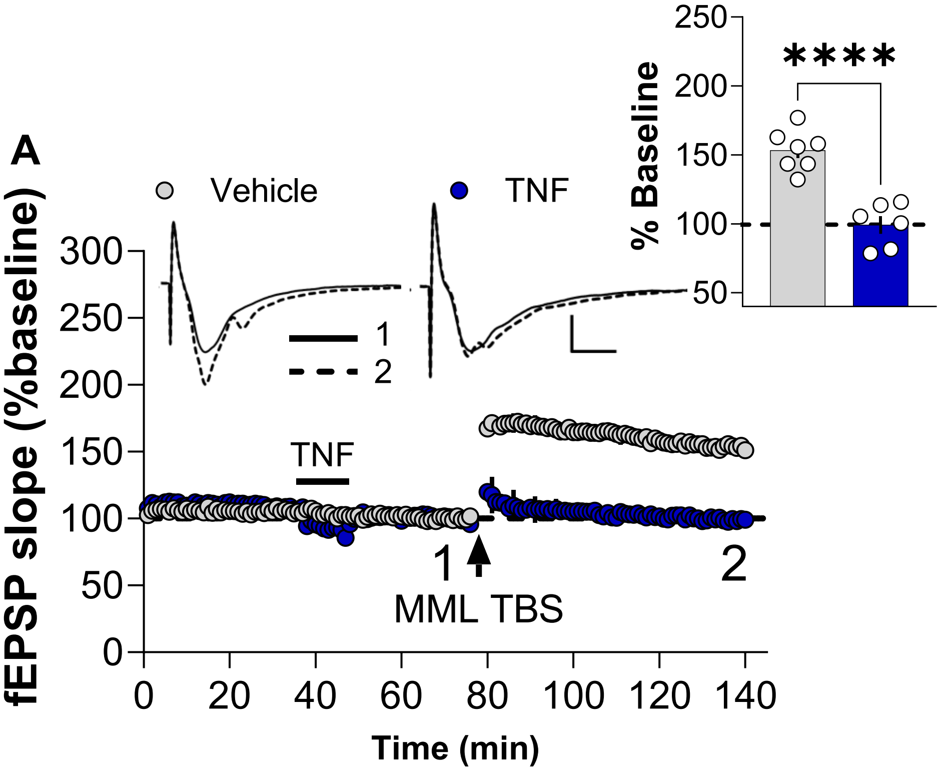


**Figure S12: TNF priming-mediated MML LTP inhibition and transregional metaplasticity**

**(A)** Bath application of 1.18 nM TNF for 10 min significantly inhibited MML LTP compared to vehicle primed controls (Vehicle Control: 153.3 ± 5.6%, $n = 7$; TNF Primed: 99.2 ± 6.5%, $n = 6$, $t_{(11)} = 6.3$, $p < 0.0001$). ***Inset:*** Representative waveforms are an average of 10 synaptic responses prior to LTP induction (**1**) and at the conclusion of the experiment (**2**). Bar graph summarizes LTP recorded from individual slices. Scale bars: 0.5 mV, 5 ms. Arrow indicates the timing of MML LTP induction. All data presented as mean ± SEM; ****, $p < 0.0001$.

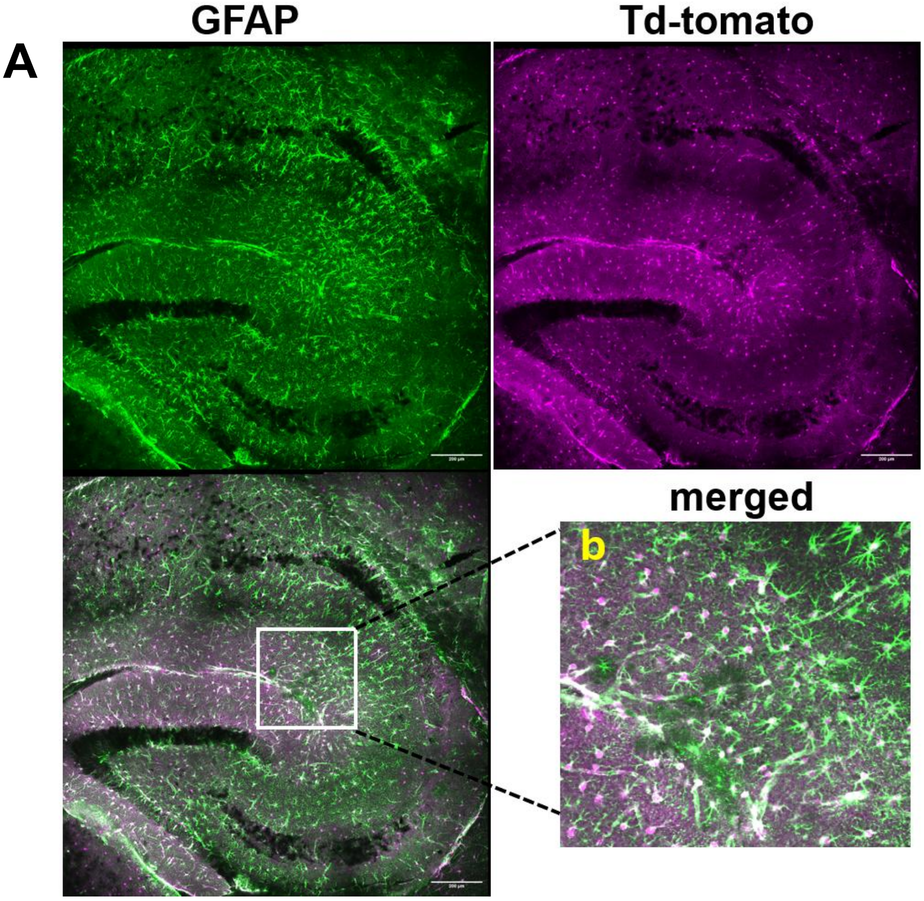


**Figure S13: Astrocytic-specific TNF knockout confirmation Td-tomato reporter for experiments described in (Fig. 4E,F).**

**(A)** Representative confocal images confirming robust, astrocytic-specific expression of the Td-tomato reporter. The expression of Td-tomato indicates successful tamoxifen-induced Cre-mediated recombination, leading to the conditional knockout of the TNF in astrocytes, which were used for experiments described in **Fig. 4E,F**. Colocalization of the magenta Td-tomato fluorescence with green GFAP immunoreactivity confirms the cell-type specificity of the knockout. Image (**b**) shows a magnified view of the boxed area in the merged panel. The slices were processed for immunofluorescence following electrophysiology recordings.

Method summary: Forty-micron-thin cryosections were fixed in 4% PFA. Antigen retrieval was performed in 10 mM sodium citrate buffer (pH 6.5) at 80 °C for one hour. Non-specific binding was blocked using 3% normal goat serum in 0.1M phosphate buffer containing 0.1% Triton X (PBTX, pH 8.5) for one hour. Sections were incubated overnight at 4°C with primary antibody against GFAP (1:1000, Chicken anti-GFAP (PA1-1004, Invitrogen), followed by incubation with secondary antibody (1:500, anti-chicken 488 (A-11039, Invitrogen) at room temperature. The colocalization of Td-tomato fluorescence (recombination reporter) with GFAP immunolabeling (astrocyte marker) validates the specific astrocytic knockout of TNF. Sections were imaged using a Nikon C2+ Confocal Microscope with a 20x objective. Scale bar: 200 μM.

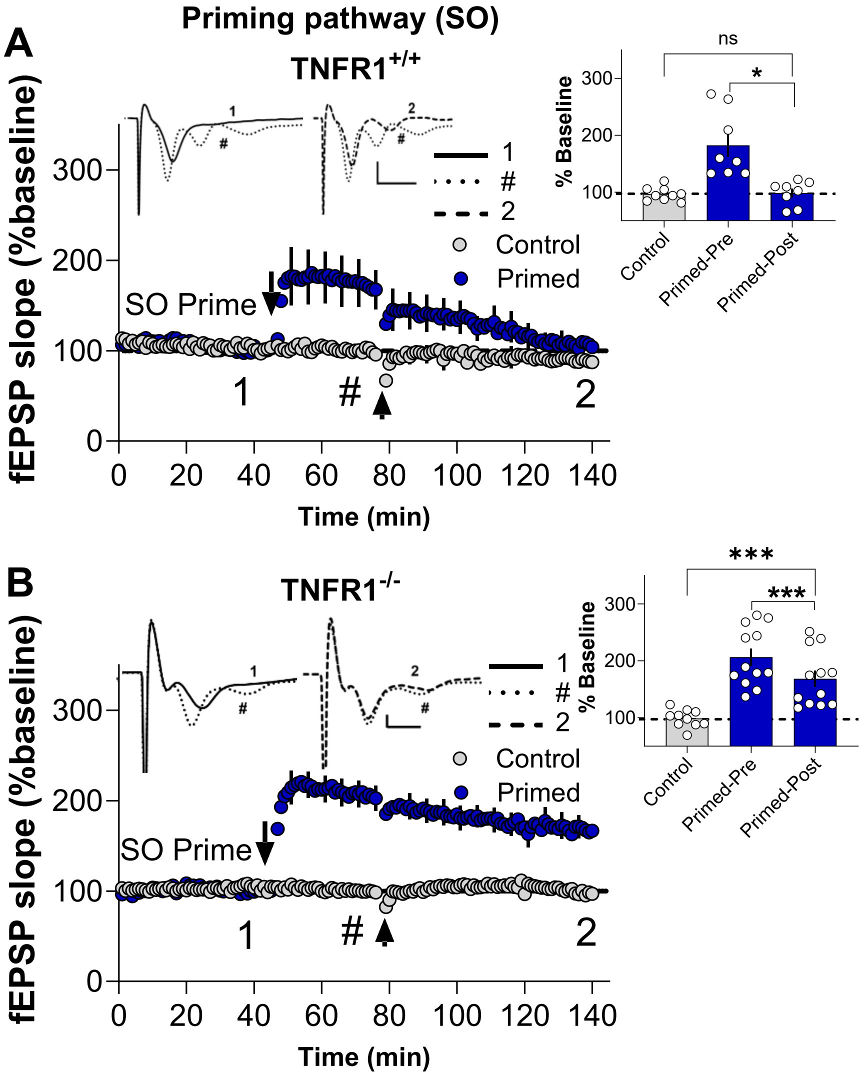


**Figure S14: SO LTP maintenance in TNFR1$^{+/+}$ and TNFR1$^{-/-}$mice from experiments described in (Fig.5 C,D)**

**(A,B)** Depotentiation of SO LTP requires TNFR1 receptors. SO priming resulted in substantial homosynaptic potentiation as measured at 30 min post-priming, both in **(A)** TNFR1$^{+/+}$ mice (Primed-Pre (#): 182.5 ± 20.7 %, $n$ = 7) and in (**B**) TNFR1$^{-/-}$ mice (Primed-Pre (#): 206.3 ± 15.0%, $n$ = 12). (**A**)TNFR1$^{+/+}$ mice showed a complete decay in SO LTP (Control: 96.5 ± 4%, $n$ = 9; Primed-Post: 98.6 ± 7.6%, $n$ = 8, $t_{(15)}$ = 0.25, $p$ = 0.81). (**B**) In the TNFR1$^{-/-}$ mice, the depotentiation of SO LTP was not observed when compared to non-primed controls (Control: 98.87 ± 4.9%, $n$ = 10; Primed-Post: 168.5 ± 14.3%, $n$ = 12, $t_{(20)}$ =14.3, $p$ = 0.0004). The sustained homosynaptic LTP observed in the TNFR1$^{-/-}$ mice suggested that the depotentiation of SO LTP in WT mice was mediated via TNFR1, as activated by the conditioning stimulus in MML. ***Inset:*** Representative waveforms are an average of 10 synaptic responses prior to SO LTP induction (**1**), 30 min post priming (**#**), and at the conclusion of the experiment (**2**). Bar graphs summarise the LTP across individual slices. Scale bars: 0.5 mV, 2 ms (**A-B**). Arrows indicate the timing of SO priming and MML LTP induction, and (**#**) indicates measurement of SO LTP 30 min post-priming or immediately before delivery of MML TBS (Primed-Pre). All data presented as mean ± SEM; *ns*, $p$ > 0.05; ∗, $p$ < 0.05; ∗∗∗, $p$ < 0.001.

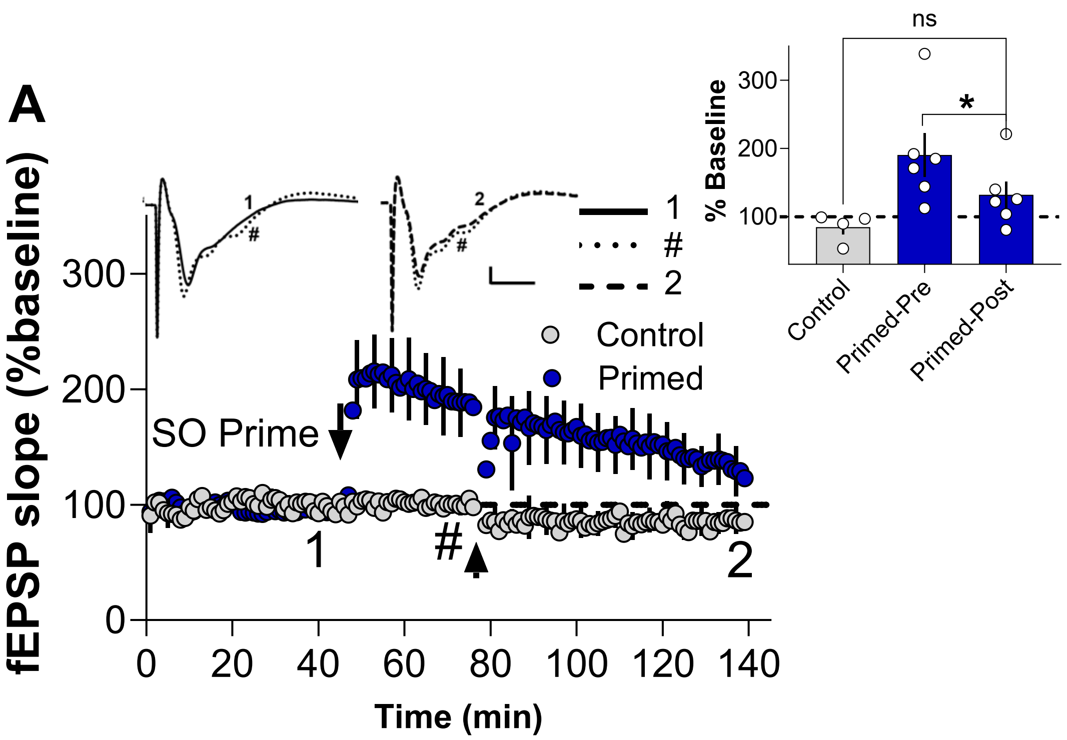


**Figure S15: Effect of Xpro1595 on priming-induced SO LTP maintenance, from experiments described in (Fig. 5H)**

(**A**) Xpro1595 treatment 12-16 hours before experimental recording did not affect homosynaptic LTP in SO as measured 30 min post-priming (Primed-Pre (#): 190.4 ± 32%, *n = 6, paired* $t_{(5)} = 4.0$, $p = 0.0108$ ) and decreased over the course of the experiment (Control: 84.59 ± 10.71%, *n = 4*; Primed-Post: 132.1 ± 19.6%, *n* = 6, $t_{(8)} = 1.83$, $p = 0.1045$). ***Inset:*** Representative waveforms are an average of 10 synaptic responses prior to SO LTP induction (1), 30 min post priming (**#**), and at the conclusion of the experiment (**2**). Bar graphs summarize the LTP across individual slices. Scale bars: 0.2 mV, 5 ms. Arrows indicate the timing of SO priming and MML LTP induction, and (**#**) indicates measurement of SO LTP 30 min post-priming or immediately before delivery of MML TBS (Primed-Pre). All data presented as mean ± SEM; *ns*, $p > 0.05$; $*$, $p < 0.05$.

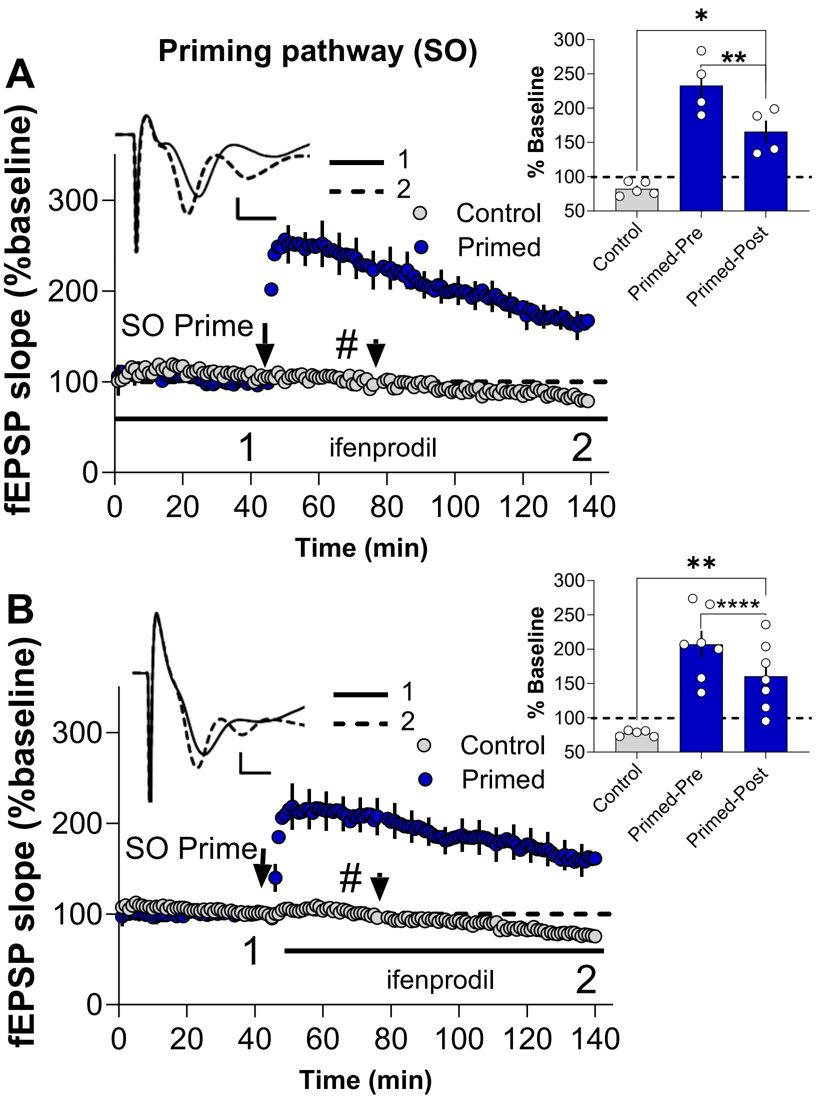


**Figure S16: Effect of the GluN2B antagonist ifenprodil on the priming-induced LTP in SO, from experiments described in (Fig. 6A,B).**

**(A)** SO priming in the presence of bath-applied 0.8 μM ifenprodil resulted in substantial homosynaptic LTP as measured at the end of 90 min (Control: 82.35 ± 4.4%; *n* = 5, Primed- Primed-Post: 165.5 ± 16.5%, *n* = 4, $t_{(7)}$ = 5.4, *p* = 0.0010), but significantly decayed over the ensuing 90 min as compared to 30 min post priming (Primed-Pre (#): 232.9 ± 2%, *n* = 4, *paired* $t_{(3)}$ = 6.2, *p* = 0.0084). **(B)** Adding ifenprodil just after SO priming gave similar results (Control: 77.46 ± 1.9%; *n* = 5, Primed- Post: 160.8 ± 18.7%, *n* = 7, $t_{(10)}$ = 3.7, *p* = 0.0041) that once again significantly decayed over time as measured 30 min post-priming (Primed-Pre (#): 207.3 ± 18.7%, *n* = 7, *paired* $t_{(6)}$ = 9.2, *p* < 0.0001). ***Inset:*** Representative waveforms are an average of 10 synaptic responses prior to LTP induction (**1**) and at the conclusion of the experiment (**2**). Bar graphs summarize the change in responses in the presence of ifenprodil. Scale bars: 0.5 mV, 2 ms (**A-B**). Arrows indicate the timing of SO priming and MML LTP induction, and (**#**) indicates measurement of SO LTP 30 min post-priming or immediately before delivery of MML TBS (Primed-Pre). All data presented as mean ± SEM; ns, *p* > 0.05; **, *p* < 0.01; ****, *p* < 0.0001.

**SI SV1: Enhanced astrocytic $Ca^{2+}$ activity in the DG molecular layer following SO Priming.**

Representative imaging of astrocyte $Ca^{2+}$ dynamics in the molecular layer of DG. The video depicts astrocytic GCaMP6f fluorescence following SO electrical priming stimulation.

**SI SV2: Astrocytic $Ca^{2+}$ activity during SO priming with mGluR and mAChR antagonists and post-washout.**

Representative imaging of astrocyte $Ca^{2+}$ dynamics in DG molecular layer in response to SO electrical priming in the presence of LY3020371 (20 μM) and pirenzepine (20 μM).
